# Excessive E2F transcription in single cancer cells precludes transient cell cycle exit after DNA damage

**DOI:** 10.1101/2020.03.19.998674

**Authors:** Hendrika A. Segeren, Lotte M. van Rijnberk, Eva Moreno, Frank M. Riemers, Ruixue Yuan, Richard Wubbolts, Alain de Bruin, Bart Westendorp

## Abstract

E2F transcription factors control the expression of cell cycle genes. Cancers often demonstrate enhanced E2F target gene expression, which can be explained by increased percentages of replicating cells. However, we now demonstrate in human cancer biopsies that individual neoplastic cells display abnormally high levels of E2F-dependent transcription. To mimic this situation, we deleted the atypical E2F repressors (E2F7/8) in untransformed cells. Individual cells with elevated E2F-activity during S/G2-phase failed to exit the cell cycle after DNA damage and underwent mitosis. In contrast, wild type cells completed S-phase and then exit the cell cycle by activating the APC/C^Cdh1^ via repression of the E2F-target Emi1. Strikingly, many arrested wildtype cells could eventually inactivate APC/C^Cdh1^ to execute a second round of DNA replication and mitosis, thereby becoming tetraploid. Cells with elevated E2F-transcription fail to exit the cell cycle after DNA damage which potentially causes genomic instability, promotes malignant progression and reduces drug sensitivity.

## Introduction

The decision to commit to a new round of S-phase is a pivotal step in the cell cycle. This step is irreversibly enforced by activation of E2F transcription factors and inactivation of the Anaphase-promoting complex/cyclosome (APC/C) E3 ligase complex (Cappell, Chung, et al, 2016). In presence of sufficient growth factor signaling, cyclins become expressed, which bind and activate cyclin-dependent kinases. This leads to the phosphorylation of the Retinoblastoma protein (RB1). RB phosphorylation releases its inhibitory effect on activator E2Fs (E2F1-3). These transcription factors then activate expression of a large network of genes involved in S-phase entry and progression. These E2F target genes include cyclin E and Emi1, which inactivate the APC/C^Cdh1^ to allow rapid accumulation of multiple proteins that are essential for S-phase progression. As S-phase proceeds, E2F-dependent transcription is silenced again, which is then mediated by atypical repressor E2Fs (Bertoli, Skotheim, and de Bruin, 2013; Kent, and Leone, 2019).

It is widely recognized that E2F-dependent transcription is almost always increased in neoplastic versus normal tissues. In fact, high expression of this transcription program correlate strongly with poor prognosis in various types of cancer (Kent, Rakijas, et al, 2016; Lan, Bian, et al, 2018). However, it is to date unclear if the increase in E2F-target gene expression in cancer is only simply explained by the fact that the number of cycling cells in a neoplasm is increased or if individual cells have also increased E2F-target gene expression. In cells that already have entered S-phase, heterogeneity in E2F-dependent transcription may have profound effects on cell cycle fates, especially under conditions causing replication stress and DNA damage since E2F targets include DNA replication and repair genes (Bertoli, Herlihy, et al, 2016). Cells are vulnerable to DNA damage during S-phase, and under conditions of genotoxic stress, cells must decide to either repair the damage and proceed to undergo mitosis, or to arrest and exit the cell cycle (Gire, and Dulic, 2015). The machinery underlying this decision point must also show switch-like behavior, because the decision to either arrest or continue the cell cycle is binary. However, in contrast to the G1/S transition, the pathways underlying the switch-like behavior of cells enforcing a cell cycle arrest are far less well-studied than the G1/S transition.

We propose that the combined action of factors controlling E2F-dependent transcription once the cell cycle is ongoing determines the decision to stop or continue cycling during genotoxic stress. These factors include expression of activator and repressor E2Fs, cyclin-dependent kinases (CDKs), ubiquitin ligases and CDK-inhibitors, many of which can be disturbed in cancer cells. To address this idea, we used single-cell analysis techniques to study heterogeneity in E2F-dependent expression in neoplastic versus normal human cells. In addition, we studied how manipulation of E2F-dependent transcription during S/G2-phase affected cell cycle decision-making after DNA damage.

Here, we show that E2F-dependent transcription is elevated at a single cell level in cycling human cancer cells from cancer biopsies. Furthermore, we found that deregulation of E2F-dependent transcription during S/G2-phase in non-transformed cells had profound consequences on cell cycle fates in response to DNA-damage. Specifically, we found that the atypical E2F repressor proteins E2F7/8 are of critical importance in enforcing a long term arrest of G2 cells after genotoxic stress. We characterized this as a 4N-G1 state, because a large subset of arrested cells could eventually re-enter the cell cycle to undergo another round of S-phase and subsequent mitosis, leading to tetraploidy. In contrast, the E2F7/8-mutant cells showed a severe defect in undergoing such a 4N-G1 arrest and tetraploidization, and instead directly progressed to undergo unscheduled mitosis. Knockdown experiments showed that repression of the E2F-target gene Emi1 - which inhibits the APC/C^Cdh1^ complex - plays a critical role herein. Finally we demonstrate that E2F7/8 repressors cooperate with P53 and its target gene P21 to enforce the 4N-G1 arrest. Hence, the combined action of multiple mechanisms affecting expression of E2F target genes -in particular Emi1-underlies the switch-like decision of cells to arrest in G2 or proceed to mitosis. This decision is pivotal to control genomic integrity in human cells.

## Results

### Abnormally high E2F-dependent transcription in individual cancer cells detected in human biopsies

We leveraged public available single cell RNA sequencing data to answer the question if E2F-dependent transcription is elevated in cancer at the single cell level. First we analyzed a dataset containing almost 6000 cells from 18 head and neck squamous cell carcinoma (HNSCC) patients (Puram, Tirosh, et al, 2017). As a proxy for overall E2F transcription, we computed for each cell the average z-scores of a set of 80 different E2F target genes (Figure 1A). We previously verified these 80 genes as E2F targets using ChIP-sequencing, motif analysis, and microarrays (Westendorp, Mokry, et al, 2012). Dimensionality reduction using t-distributed stochastic neighbor embedding (tSNE) revealed well-separated clusters of different stromal cell types and three main clusters of malignant cells (Figure 1B, upper panel). The three tumor cell clusters clearly contained more cells with high E2F-scores when we projected these scores on the tSNE maps (Figure 1B, lower panel). Boxplots also demonstrated that E2F scores were clearly elevated in HNSCC cancer cells versus all non-tumor cells (Figure 1C). Plotting RNA counts of individual E2F target genes consistently showed the same pattern (Figure 1C – insets), suggesting that indeed single cancer cells exhibit enhanced E2F-dependent transcription.

**Figure 1.**
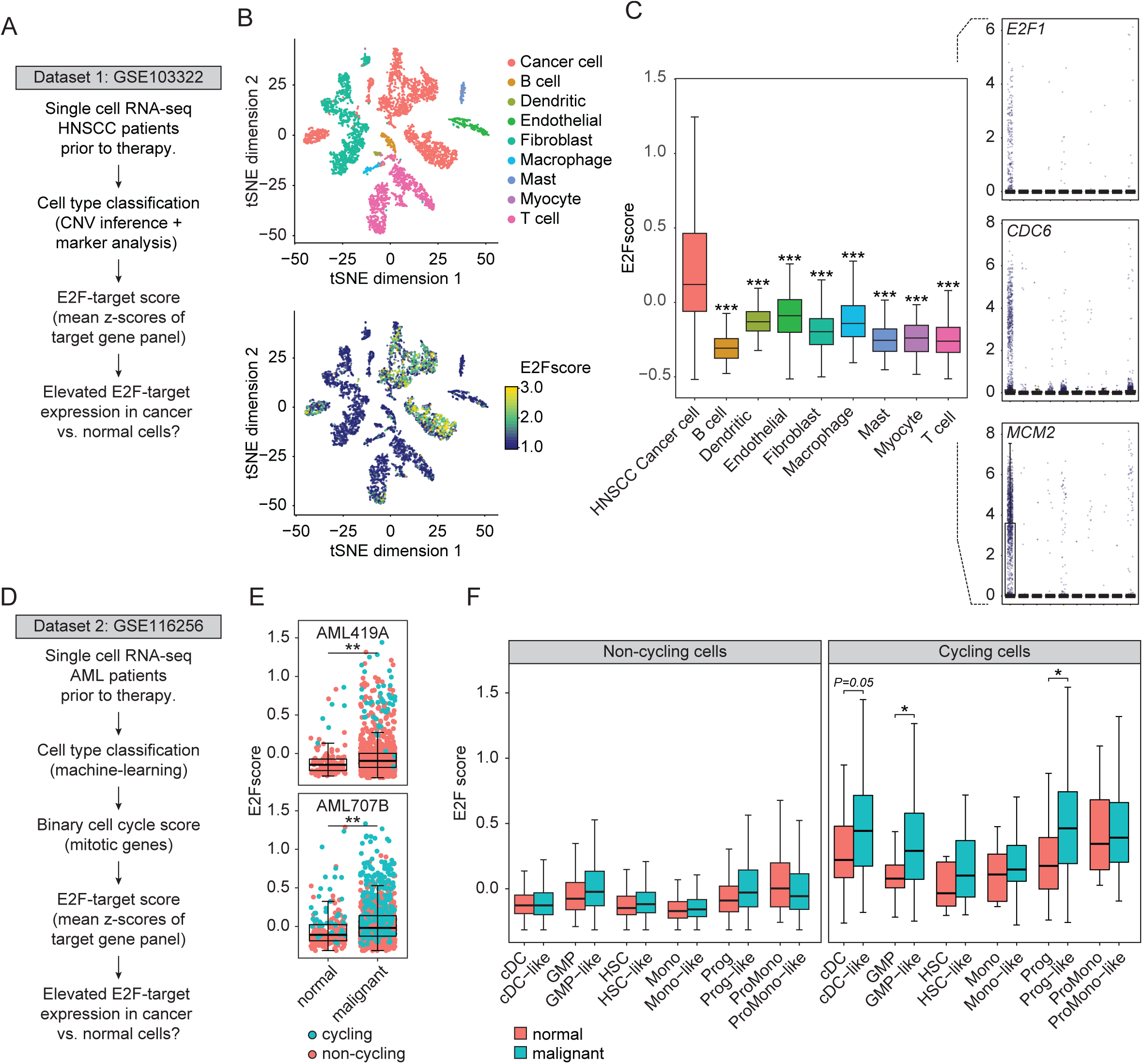
Elevated E2F-dependent transcription in malignant versus non-transformed single cells. **A** Schematic overview of the workflow to analyze E2F-target gene expression in single cells from patients with head and neck squamous cell carcinoma (HNSCC). *** *P*<0.0001 as indicated, Kruskall-Wallis test followed by Dunnet’s pairwise comparisons with post-hoc Benjamini-Hochberg correction. **B** Dimensionality reduction using t-Stochastic Neighbourhood Embedding (tSNE) shows clusters of HNSCC tumor cells and different types of stromal cells. **C** Boxplots representing E2F-target expression scores of HNSCC tumor cells and different stromal cell types. Insets show dot plots with corrected mRNA counts of three representative E2F-target genes included in the E2F score calculation. **D** Schematic overview of the workflow to analyze E2F-target gene expression in single cells from patients with acute myeloid leukemia (AML). **E** E2F target gene expression scores in malignant versus non-malignant cells from two individual AML patients. Cells are color-labeled according to binary cell cycle classification scores, as determined in the original publication (van Galen, Hovestadt, et al, 2019) **F** Boxplots representing E2F-target expression scores of malignant versus normal cells from AML patients, grouped according to cell-type classification. cDC: conventional dendritic cell, GMP: granulocyte-macrophage progenitor, HSC: hematopoietic stem cell, Mono: monocyte, Prog: progenitor, ProMono: promonocyte. The addition “-like” refers to the best matching normal cell type identity in malignant cells * *P*<0.05 as indicated, Wilcoxon paired-rank tests with post-hoc Benjamini-Hochberg correction.

A limitation of this dataset is that HNSCC cells could not directly be compared to their non-transformed counterparts (i.e. normal epithelium). We could only compare tumor to various stromal cell types. Ideally we would want to compare E2F-dependent transcription in cycling cancer cells and normal cells within the same tissue type. Therefore, we analyzed a dataset from acute myeloid leukemia (AML) patients (van Galen, Hovestadt, et al, 2019). AML is a rapidly proliferating cancer, but several normal cell types in the blood proliferate rapidly as well. Van Galen and coworkers developed a machine-learning algorithm to classify the extent to which AML cells resemble normal white blood cell types, allowing us to compare E2F scores in malignant cells and their non-transformed counterparts (Figure 1D). In addition, they developed a scoring method to separate cycling from non-cycling cells using a gene signature of mitotic genes (van Galen, Hovestadt, et al, 2019). First, we confirmed that E2F scores were overall strongly elevated in neoplastic cells versus non-transformed cell types almost every individual patient, without taking into consideration the specific cell type (Figure 1E, S1). We then compared the E2F scores between specific normal white blood cell types and their malignant counterparts, in cycling and non-cycling cells separately. Importantly, tSNE analysis revealed that only differentiated cell types, such as B cells, T cells, and natural killer cells were separated from malignant cells, in contrast to more undifferentiated cell types (Figure S2A-C). This indicates the overall resemblance in gene expression between malignant cells and their untransformed counterparts.

Interestingly, we observed large variation in E2F scores between the different malignant cell types, suggesting a large heterogeneity in E2F-dependent transcription in cancer cells (Figure 1F). When comparing malignant cell types to their non-transformed counterparts, we observed that E2F scores were elevated in cancer cells. In particular malignant cells resembling granulocyte-macrophage progenitors (GMPs), and progenitors (Progs) showed a statistically significant enhancement of E2F-dependent transcription compared to their non-transformed cycling counterparts (Figure 1F). Together, these data strongly suggest that E2F-dependent transcription is highly heterogeneous between different types of cycling cells, and in particular elevated in cancer cells. As E2F-target genes include genes involved in DNA-repair and cell cycle progression, variation can have profound effects on cell cycle fates, beyond overriding the G1/S checkpoint. However it is challenging to study this further in patient-derived material, because cell cycle stage could only be inferred from transcriptomic information. Furthermore, it is impossible to dissect the direct consequences of E2F de-repression during S/G2 phase, and the indirect effects of overriding the G1/S checkpoint. Therefore, we decided to analyze the effects of unrestrained E2F-dependent transcription on DNA damage responses at single cell level in a more controlled setting, i.e. in cells expressing high levels of E2F transcription during S- and G2 phase. To this end, we used RPE cells carrying the FUCCI reporter system, in which we had deleted E2F7 and E2F8 using CRISPR/Cas9 (Figure S3A-B). We made double mutants (hereafter referred to as *E2F7/8*^*KO*^ cells), because substantial functional redundancy exists between these two highly homologous family members (Li, Ran, et al, 2008; Thurlings, Martinez-Lopez, et al, 2017).

### Deregulated E2F-dependent transcription during S/G2-phase causes cell cycle fate defects

Since E2F7/8 are only present during S/G2-phase of the cell cycle (Boekhout, Yuan, et al, 2016), we first wanted to establish that *E2F/8*^*KO*^ FUCCI cells specifically show target gene deregulation during these phases. The FUCCI system allows to distinguish G1-cells from S- and G2-cells via the alternating degradation of Azami Green (mAG-) tagged truncated Geminin and degradation of Kusabira Orange (mKO-) tagged truncated CDT1 (Sakaue-Sawano, Kurokawa, et al, 2008). Therefore, we sorted cells according to expression levels of the FUCCI reporters, and then quantified the expression of multiple E2F target genes in control and *E2F7/8*^*KO*^ cells. We observed that E2F7/8 loss only enhanced the expression of target genes when cells were in S- or G2-phase (Fig 2A). This suggests that *E2F7/8*^*KO*^ cells still have an intact G1/S checkpoint. In fact, live cell imaging showed that *E2F7/8*^*KO*^ G1-cells were less likely to enter S-phase under unperturbed conditions than control cells (Figure 2B). This was also reflected by lower proliferation rates during unperturbed conditions (Figure S3D).

**Figure 2.**
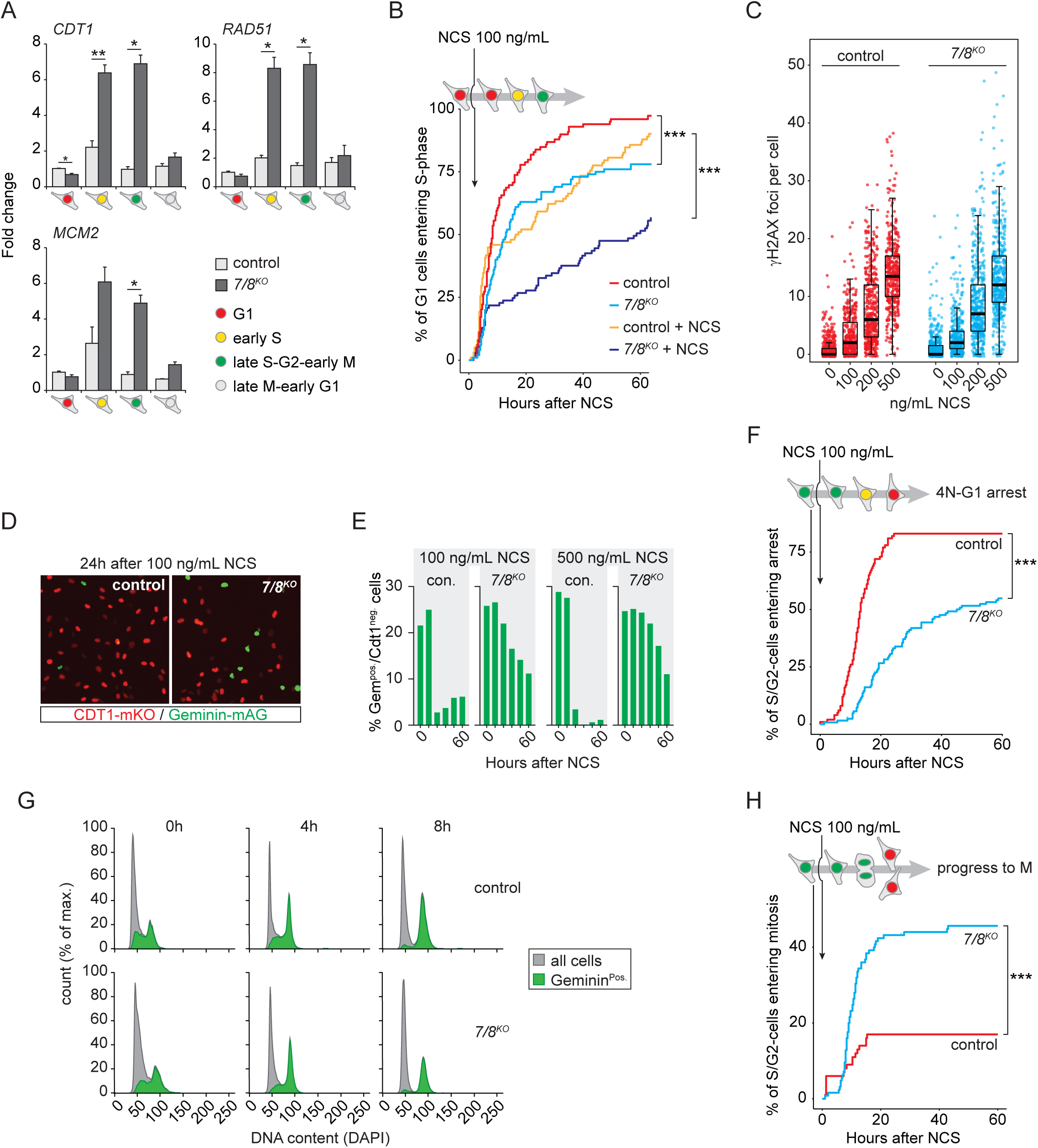
Upregulation of E2F-target gene expression during S/G2-phase by E2F7/8 deletion results in aberrant S/G2-phase DNA damage response. **A** Quantitative PCR on FACS-sorted FUCCI-expressing cells, showing cell cycle-dependent deregulation of E2F target genes in *E2F7/8*^*KO*^ cells. Bar represents mean ± s.e.m of in total three experiments. * *P*<0.05, ** *P*<0.005. **B** Time until S-phase entry in control and *E2F7/8*^*KO*^ RPE-FUCCI cells under unperturbed conditions or after addition of 100 ng/mL neocarzinostatin (NCS), quantified as cumulative entry of G1 cells into S-phase during live cell imaging. S-phase entry was determined by appearance of geminin. Data from two independent cell clones were pooled, n=100 cells in total, 50 per clone. **C** Quantification of γH2AX foci, 8 hours after a single dose of neocarzinostatin (NCS). Each dot represents a nucleus. **D** Snapshot of RPE-FUCCI cells, taken 24 hours after treating cells with 100 ng/mL NCS. **E** Quantification of percentage Geminin^pos^/CDT1^neg^ cells at 12-hour intervals after NCS treatment. In each condition, at least 500 cells were quantified. **F** Cumulative numbers of cells re-activating the APC/C ^Cdh1^ during G2 in response to NCS. Per condition fates of 100 S/G2 cells from two independent control and *E2F7/8*^*KO*^ clones were determined during 60 hours of live cell imaging. **G** Flow cytometric analysis of DNA content in RPE-FUCCI cells in response to 100 ng/mL NCS (4 and 8 hours). Green histograms represent Geminin^pos^/CDT1^neg^ cells. **H** Quantification of S/G2 cells that underwent mitosis without interrupting G2 phase, observed by live cell imaging. Per condition, at least 100 cells from 2 different CRISPR clones were followed. Statistical analysis in B, F, and H was performed with Log Rank analysis.

We then treated cells with the radiomimetic drug NCS, which causes double-stranded DNA breaks (DSBs) irrespective of cell cycle phase (Chao, Poovey, et al, 2017). Furthermore NCS is highly unstable, which allows cells to recover without the need to wash the drug away during live cell imaging experiments. When treated with 100 ng/mL NCS, the *E2F7/8*^*KO*^ G1 cells were significantly less likely to enter S-phase during the imaging period (Figure 2B, S3C). These observations could be explained by the notion that E2F-deregulation in the *E2F7/8*^*KO*^ cells causes mild replication stress-induced DNA damage, which cannot be completely resolved prior to mitosis. Unresolved endogenous replication stress in the mother cells can cause daughter cells to enter quiescence (Arora, Moser, et al, 2017). To test this hypothesis, we transduced the FUCCI expressing cells with a lentiviral construct carrying mTurquoise-tagged 53BP1. Presence of 53BP1 foci during G1 are indicative of replication stress-induced DNA damage in the previous cell cycle. And indeed, we found that a significantly increased percentage of *E2F7/8*^*KO*^ nuclei in G1 contained 53BP1 foci (Figure S3E).

The modest increase in replication stress-induced 53BP1 foci under unperturbed conditions led us to investigate if uncontrolled E2F-dependent transcription during S-phase also results in more exogenous DNA damage. First, we quantified DNA damage with immunofluorescence staining of γH2AX. NCS dose-dependently induced similar levels of DNA damage in control and *E2F7/8*^*KO*^ cells (Figure 2C). Although a low dose (100 ng/mL) of NCS only caused modest DNA damage, it was sufficient to strongly inhibit proliferation of both control and *E2F7/8*^*KO*^ cells over a period of 60 hours of imaging (Figure S3D). Despite similar levels of DNA damage, *E2F7/8*^*KO*^ cells showed a strong increase in the percentage of geminin-positive / CDT1-negative (Geminin^pos.^/CDT1^neg.^) cells after NCS treatment, indicating an enhanced number of cycling cells despite DNA damage (Figure 2D). We found that the percentage of Geminin^pos.^/CDT1^neg.^ cells was markedly increased in *E2F7/8*^*KO*^ compared to control cells between 1 and 3 days after NCS, both after a moderate and a high dose of NCS (respectively 100 and 500 ng/mL, Figure 2E). According to the expected cell-cycle dependent degradation of the FUCCI reporters, these are cells in mid-to-late S phase or G2. Therefore, we followed the fates of cycling -i.e. Geminin^pos.^/CDT1^neg.^-cells after addition of NCS. Indeed, we saw that within 24 hours the far majority of the control RPE cells in S/G2-phase responded to NCS by exiting the cell cycle and entering a G1-like state, as seen by loss of geminin and reappearance of CDT1 without mitosis (Figure 2F, SUPP MOVIE). Disappearance of the Geminin-mAG signal is caused by activation of the APC/C^Cdh1^ complex. This complex is active during late mitosis and G1, but can also be re-activated in response to DNA damage to exit the ongoing cell cycle(Bassermann, Frescas, et al, 2008). This arrest was severely delayed in *E2F7/8*^*KO*^ cells (Figure 2F), explaining the elevated level of Geminin^pos.^/CDT1^neg^ cells compared to control cells. We then asked if the re-activation of the APC/C^Cdh1^ already happened during S-phase, or whether cells first completed S-phase to arrest in G2 in response to NCS. To answer this question, we analyzed their DNA content with flow cytometry. This analysis showed that by 8 hours after addition of NCS, virtually all Geminin^pos.^/CDT1^neg.^ cells had a 4N DNA content (Figure 2G). This shows that the cells first finish S-phase, before entering the G1-like state.. We will therefore refer to this cell cycle exit as a 4N-G1 arrest Further cell fate analysis of the Geminin^pos.^/CDT1^neg.^ cells showed that more than twice as many Geminin^pos.^/CDT1^neg.^ *E2F7/8*^*KO*^ as control cells did not arrest in response to 100 ng/mL NCS, and proceeded to unscheduled mitosis (Figure 2H). Together, these results show that cells in S- or G2-phase respond to DNA double stranded breaks with a 4N-G1-like arrest in an E2F7/8-dependent manner.

### The G1-like state in NCS-treated cells is highly similar to a normal G1, and is essential for DNA damage recovery

Hitherto, we defined the G1-like state by only the loss of geminin and appearance of CDT1. To gain further insight into this biological state we performed single cell RNA-sequencing on control and *E2F7/8*^*KO*^ cells, 24 and 48 hours after treatment with NCS. To this end, we used an automated RNA-sequencing platform (Muraro, Dharmadhikari, et al, 2016). However, instead of capturing the cells with FACS-sorting, we isolated and captured cells with a VyCAP needle puncher (Figure 3A). This system is designed to take fluorescence images of up to 6400 individual cells, from which cells of interest can be automatically selected and isolated for single cell genomics and transcriptomics (Stevens, Oomens, et al, 2018). Using an ultrafine needle mounted to a motorized z-stage, cells are punched from the chip into a microwell plate (Figure 3B). This procedure allowed us to perform RNA-sequencing on cells with a known FUCCI-fluorescence status. The fluorescence signals of the punched cells showed that all cell cycle phases were captured from both the untreated control and *E2F7/8*^*KO*^ cell lines (Figure 3C). However, treatment with an intermediate dose of 200 ng/mL NCS showed a marked increase in the number of quiescent or G1-like (Geminin^neg^/CDT1^pos^) cells. At 24 hours, we observed again a clear increase in the number of Geminin^pos^/CDT1^neg^ cells when comparing *E2F7/8*^*KO*^ with control cells (Figure 3C). In total 180 cells passed our rigorous quality control. We could identify four main clusters of cells, which correlated well with cell cycle phase and NCS treatment. Clusters I and II corresponded with untreated S/G2 cells and G1 cells, respectively. Clusters III and IV respectively consisted of arrested cells 24 and 48 hours after NCS treatment (Figure 3D). The expression of cell cycle genes such as *MKI67* and the E2F target gene *MCM2* showed the highest expression in cluster I, whereas stress-related genes including the P21-encoding *CDKN1A* and *FBXO32* were most highly expressed in cluster IV (48h NCS). Thus the observed 4N-G1 cell cycle arrest upon NCS treatment coincides with a transcriptional state similar to G0/G1 cells. Overall, the *E2F7/8*^*KO*^ cells and control cells were not separated by the cluster analysis, suggesting that expression profiles of non-cell cycle genes were comparable (Figure 3D). But interestingly, we observed several strongly geminin-positive NCS-treated *E2F7/8*^*KO*^ cells in clusters I and II. These cells most likely represent the previously observed S/G2-phase cells which were unable to activate APC/C^Cdh1^ and did not arrest in a G1-like state in response to NCS. Clusters I and II largely contain untreated cycling cells, suggesting that E2F7/8-mutant cells failing to degrade geminin after NCS showed S/G2-like cell cycle gene expression profiles. To further study this, we grouped the single cell gene expression profiles of the *E2F7/8*^*KO*^ cells and control cells by cell cycle status as determined by FUCCI reporters. Again, we observed upregulation of E2F target genes including *CDC6* and *MCM2* during S- and G2-phase in untreated *E2F7/8*^*KO*^ cells (Figure 3E, upper row). In contrast, FOXM1 target genes, exemplified here by *CDK1* and *PLK1*, and the general cell cycle marker MKI67 were not de-repressed in *E2F7/8*^*KO*^ cells during unperturbed G2-phase (Figure 3E, upper row). However, when examining expression of these genes after NCS treatment, expression of E2F target genes was strongly elevated in Geminin^pos.^/CDT1^neg.^ *E2F7/8*^*KO*^ cells 24 hours after NCS compared to CDT1^pos.^ cells of both genotypes (Figure 3E, bottom row). In addition, these cells showed strongly upregulated expression of mitotic B-Myb/FOXM1-target genes including *CDK1* and *PLK1*, as well as *MKI67* (Figure 3E, bottom row). This indicates that the Geminin-positive subset of *E2F7/8*^*KO*^ cells indeed remained in a S- or G2-like state. We did not observe these Geminin^pos.^/CDT1^neg.^ cells after 48 hours, but we only managed to capture a low number of cells at this time point (Figure S4A).

**Figure 3.**
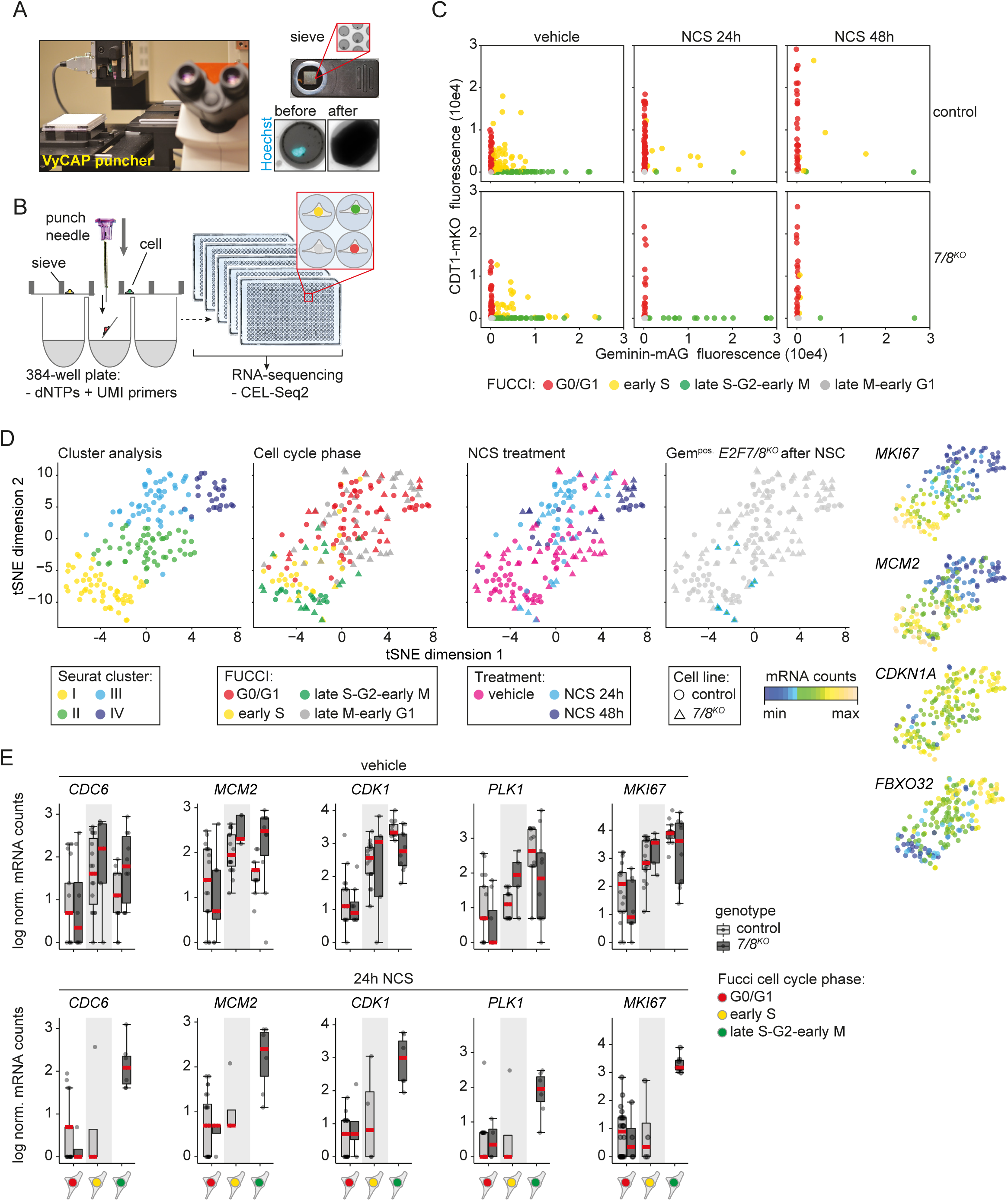
Single cell transcriptomics reveal deregulation of E2F transcription as well as the mitotic (B-Myb/FOXM1) gene expression program after E2F7/8 deletion. **A** VyCAP needle puncher system for capturing, imaging, and selection of cells for single cell sequencing. **B** Cartoon showing how the needle puncher is used to collect single cells in microwell plates for RNA-sequencing. Right panel shows how 384-well plates are processed for sequencing using CEL-Seq2; up to 5 plates can be pooled in one Illumina Nextseq run. **C** Quantification of FUCCI reporters in RPE cells captured for single cell transcriptomics. **D** tSNE plots of single cell mRNA sequencing on control and *E2F7/8*^*KO*^ cells with and without NCS treatment (200 ng/mL). Data normalization, tSNE analysis and cluster analysis were all performed using the Seurat package. **E** mRNA counts of representative E2F targets (*CDC6* and *MCM2*) and FOXM1 targets (*CDK1, PLK1*) and the general cell cycle marker *MKI67* from the single cell sequencing data, separated by treatment, cell cycle, and genotype condition.

In a parallel approach, we performed pseudo-time alignment using the Monocle algorithm (Trapnell, Cacchiarelli, et al, 2014). Monocle is a well-established tool that can use transcriptome data to order single cells according to their progression between different states, such as differentiation or cell cycle phase. This pseudo-time alignment showed that in our single cell data set cells were indeed ordered according to expression changes in cell cycle genes and genes related to cell stress (Figure S4B).

Importantly, Monocle analysis confirmed in an unsupervised manner that Geminin^pos.^/CDT1^neg.^ *E2F7/8*^*KO*^ cells after NCS treatment maintained an S/G2-like expression profile after NCS treatment (Figure S4B-C). Together, these single cell RNA-sequencing data show that the DNA damage-induced cell cycle arrest indeed resembles a G1-like gene expression profile, and that atypical E2Fs are of pivotal importance to enforce these transcriptomic changes.

### Repression of the E2F7/8 target gene FBXO5 is required to mediate the 4N-G1 arrest in response to DNA damage

Because atypical E2Fs are transcriptional repressors, we asked downregulation of which target genes are critical to enter a G1-like state after DNA damage. The G1-like state is established by re-activation of APC/C^Cdh1^, as exemplified by APC/C dependent degradation of the FUCCI reporter Geminin. A key candidate effector would be *FBXO5*, which encodes the endogenous APC/C^Cdh1^ inhibitor Emi1. Previous work showed that P53-dependent activation of P21 can activate APC/C^Cdh1^ via repression of Emi1 expression in arrested G2 cells after genotoxic stress (Lee, J., Kim, et al, 2009; Wiebusch, and Hagemeier, 2010). We previously demonstrated that Emi1 is transcriptionally regulated by E2F7 and −8 (Boekhout, Yuan, et al, 2016). *FBXO5* transcripts were only modestly increased by E2F7/8 deletion in the FACS-sorted unperturbed FUCCI cells (Figure 4A). However, the single cell RNA-sequencing data showed that FBXO5 transcripts were highly elevated in the Geminin^pos.^/CDT1^neg.^ cells lacking E2F7/8 after NCS, compared to G1-like NCS-treated control cells (Figure 4B). To further test our hypothesis, we used RNAi immediately after NCS addition to rescue the elevated expression of *FBXO5* (Emi1) in *E2F7/8*^*KO*^ cells, and then counted the presence of geminin-positive cells after 24 hours (Figure 4C). Emi1 protein was only slightly increased in unperturbed *E2F7/8*^*KO*^ cells (Figure 4D). However, we observed a huge increase in Emi1 protein in NCS-treated *E2F7/8*^*KO*^ cells compared to control cells (Figure 4E). Interestingly, a low dose of 1 nM of Emi1 siRNA was already sufficient to abolish the Emi1 overexpression and rescue the percentage of Geminin^pos^/CDT1^neg^ *E2F7/8*^*KO*^ cells to a level comparable to control cells (Figure 4F).

**Figure 4.**
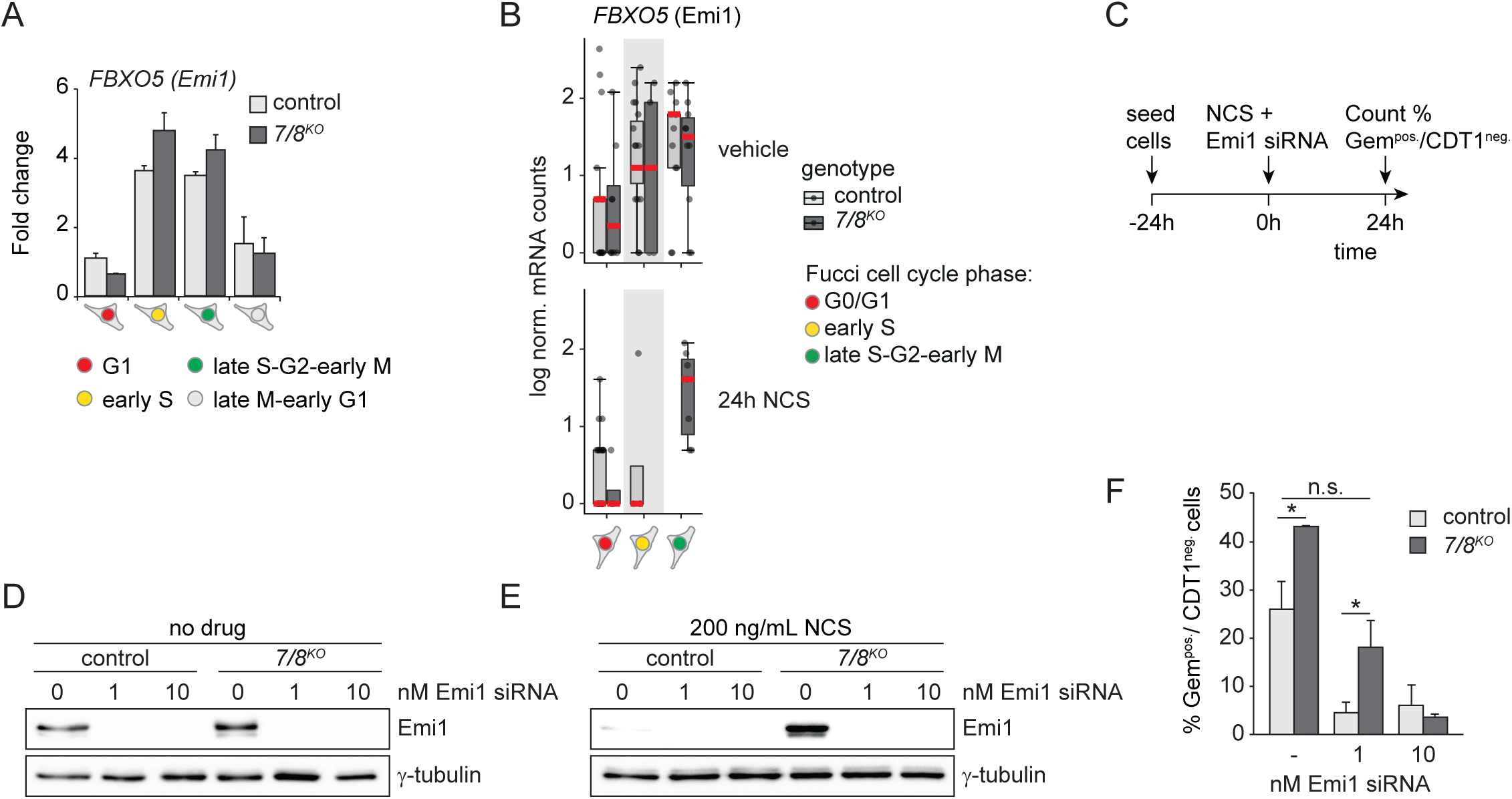
Failure of *E2F7/8*^*KO*^ cells to enter a G1-like arrest after S-phase is caused by upregulation of Emi1 expression. **A** Quantitative PCR on FACS-sorted FUCCI-expressing cells, showing expression of *FBXO5* -which encodes the APC/C^Cdh1^ inhibitor protein Emi1-in unperturbed *E2F7/8*^*KO*^ and control cells. Bar represents mean ± s.e.m of in total three experiments. **B** Transcript counts of *FBXO5* in RPE-FUCCI cells, measured with single cell RNA-sequencing. Note high levels in S/G2 *E2F7/8*^*KO*^ cells 24h after NCS treatment. **C** Experimental design of Emi1 siRNA transfections to rescue the capacity of *E2F7/8*^*KO*^ cells to enter a 4N-G1 arrest after NCS treatment. **D** Immunoblot showing expression of Emi1 in unperturbed RPE-FUCCI cells, and 24 hours after addition of siRNA against Emi1. Blots are representative examples of two independent experiments. **E** Same as in D, but now in presence of NCS. **F** Knockdown of FBXO5 immediately after NCS treatment rescues the G1-like arrest in E2F7/8 ^*KO*^ cells. Percentage of Geminin^pos.^/CDT1^neg.^ cells were counted from fluorescence microscopy snapshots of living cells, 24 hours after 200 ng/mL NCS. Bars represent the average ± s.e.m. of in total 2 experiments (two clones of cells in each experiment, n=500 cells per condition). Differences in percentages of cells per condition were statistically evaluated using Fisher’s Exact tests * *P*<0.05.

P53 and its target P21 are also pivotal in downregulation of Emi1 to maintain a cell cycle arrest in G2 after DNA damage (Lee, J., Kim, et al, 2009; Wiebusch, and Hagemeier, 2010). Other work showed that E2F7 is a direct transcriptional target gene of P53 (Aksoy, Chicas, et al, 2012; Carvajal, Hamard, et al, 2012). This raised the question to what extent E2F7/8 act in parallel with or downstream of P53 in this response. To this end, we knocked down *TP53* using siRNA and again quantified APC/C^Cdh1^ activation by counting the percentages of Geminin^pos.^/CDT1^neg.^ cells after NCS treatment (Figure 5A). RNAi treatment caused a robust knockdown of *TP53*, which resulted in strongly attenuated expression of its downstream target P21 after NCS (Figure 5B, S5A). We observed that *TP53* knockdown caused a marked increase in Geminin-positive cells after NCS treatment (Figure 5C). Interestingly, the combination of E2F7/8 deletion and P53 knockdown resulted in a slight further increase in Geminin-positive cells, suggesting that E2F7/8-dependent activation of APC/C ^Cdh1^ after DNA damage occurs mostly as a downstream effect of P53 activation (Figure 5C). In line with these observations, we found that E2F target genes in NCS-treated were upregulated by *TP53* RNAi, in particular in combination with *E2F7/8* deletion (Figure 5D).

**Figure 5.**
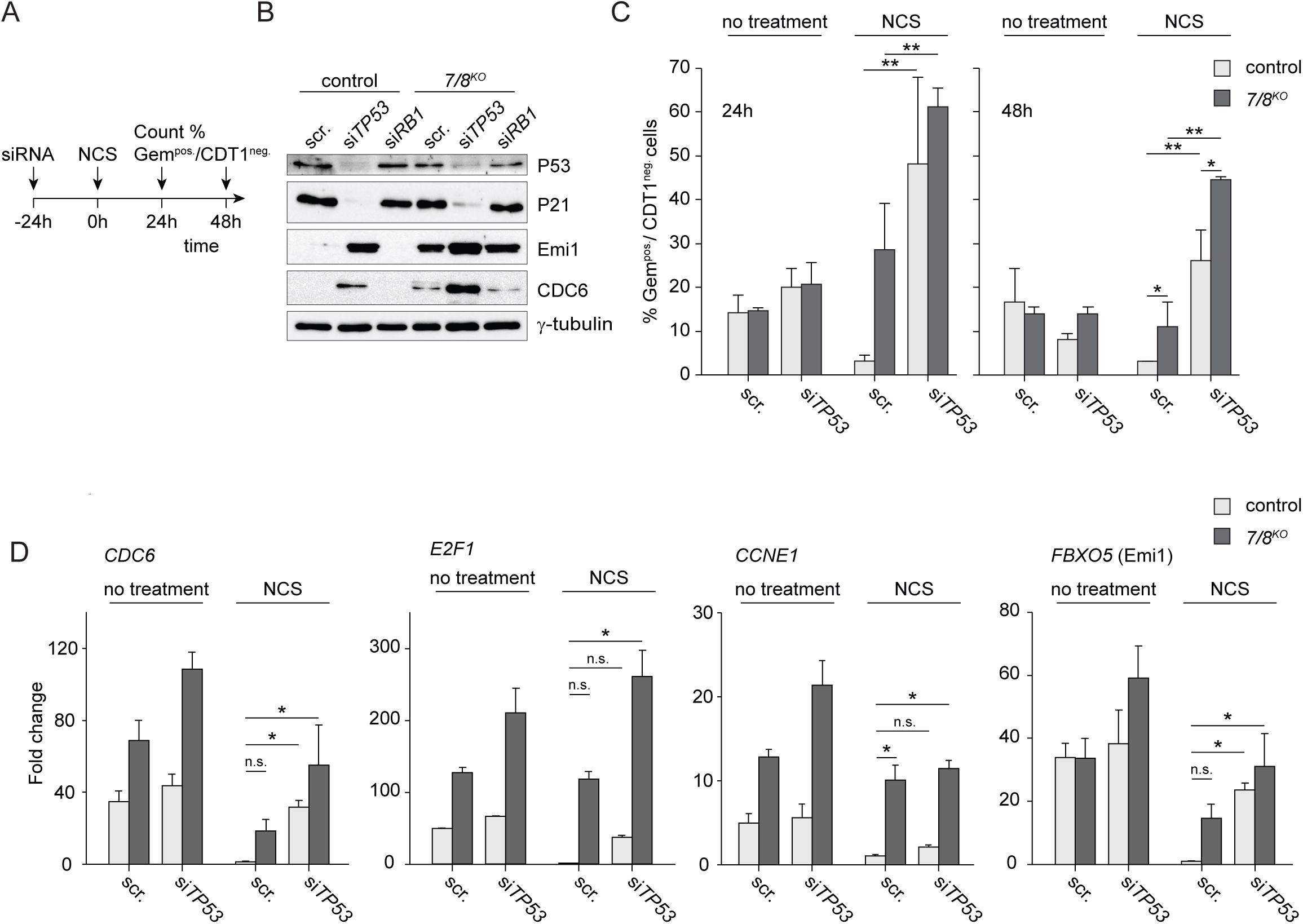
E2F7/8 and P53 cooperate to enforce a 4N-G1 arrest after DNA damage. **A** Experimental scheme of TP53 siRNA knockdown experiments to study interaction between E2F7/8 and P53 after NCS treatment. **B** Immunoblot showing expression of indicated proteins in RPE-FUCCI cells, 48 hours after addition of P53 siRNA and 24 hours after treatment with 500 ng/mL of NCS. Blots are representative examples of two independent experiments. **C** Quantification of the percentages of S/G2 (Geminin-mAG positive) cells 24 and 48 hours after 500 ng/ml NCS treatment. At least n=700 cells per condition were counted. Differences in percentages of cells per condition were statistically evaluated using Fisher’s Exact tests * *P*< 0.05, ** *P*<0.005 **D** Effects of TP53 knockdown and E2F7/8 deletion on E2F target genes in unperturbed conditions and 24 hours after NCS treatment. Bars represent mean ± s.e.m. of duplo measurements in two independent cell clones.

Importantly, these upregulated target genes included the activator E2F family member *E2F1*, the aforementioned APC/C^Cdh1^ inhibitor Emi1, and also Cyclin E1. This cyclin is required for CDK2 activation, which in turn also inhibits the APC/C to facilitate cell cycle progression, which provides an alternative route how *E2F7/8* loss prevents a 4N-G1 arrest. Previous work suggests that P21-dependent RB dephosphorylation could at least in part explain the reduced Emi1 expression to enforce the cell cycle arrest in G2 cells (Lee, J., Kim, et al, 2009). Hence, we wanted to directly compare the importance of canonical repression via RB with the importance of atypical E2Fs on E2F target gene regulation and the initiation of a 4N-G1 arrest after DNA damage. To test this, we knocked down *RB1* in the same experimental setting as TP53 (Figure 5A). Although RB1 protein and transcripts were nearly absent after siRNA treatment, we did not find a significant effect on the percentage of Geminin-positive cells after NCS (Figure S5B-D). In line with this, *RB1* knockdown caused only a minor increase in E2F target gene expression (Figure S5E). These findings suggest that RB1 does not play an important role in mediating the 4N-G1 arrest. It is possible that other pocket proteins (P107 and P130) act redundantly with RB to sustain repression of E2F target genes under DNA damaging conditions (Helmbold, Kömm, et al, 2009).

Together, these results strongly suggest that the combined action of atypical repressor E2Fs and P53 during S- and G2-phase regulates expression of E2F target genes including Emi1, to mediate APC/C^Cdh1^ re-activation and subsequent arrest of cycling cells in response to DNA damage, independently from RB1.

### Atypical E2Fs mediate DNA damage-induced endocycles

We referred to the NCS-induced cell cycle exit as a 4N-G1 arrest, but G1 implicates that these cells could escape the arrest and start a new round of S-phase. And indeed, live cell imaging revealed that a remarkably high percentage of ∼30% of the NCS-treated S/G2 cells could re-enter the cell cycle and complete mitosis after initially undergoing a 4N-G1 arrest (Figure 6A,B). However, cell cycle re-entry was virtually absent in the *E2F7/8*^*KO*^ cells (Figure 6B). Accordingly, flow cytometry analysis showed the appearance of a substantial population of cycling (i.e. Geminin-positive) cells with a 4-8N DNA content 72 hours after NCS treatment in control RPE cells, but not in *E2F7/8*^*KO*^ cells (Figure 6C-D). This tetraploidization phenomenon also led to gross mitosis defects, such as tripolar spindles. When analyzing the live cell imaging data, we observed that ∼30% of all mitoses of endocycling cells were tripolar, which can be expected to cause aneuploidy (Figure 6E). Collectively these data demonstrate that cycling cells treated with a DNA-damaging drug can escape a 4N-G1 arrest, that can eventually lead to tetraploidy and aneuploidy. Atypical repressor E2Fs are critically important in mediating this route to polyploidy.

**Figure 6.**
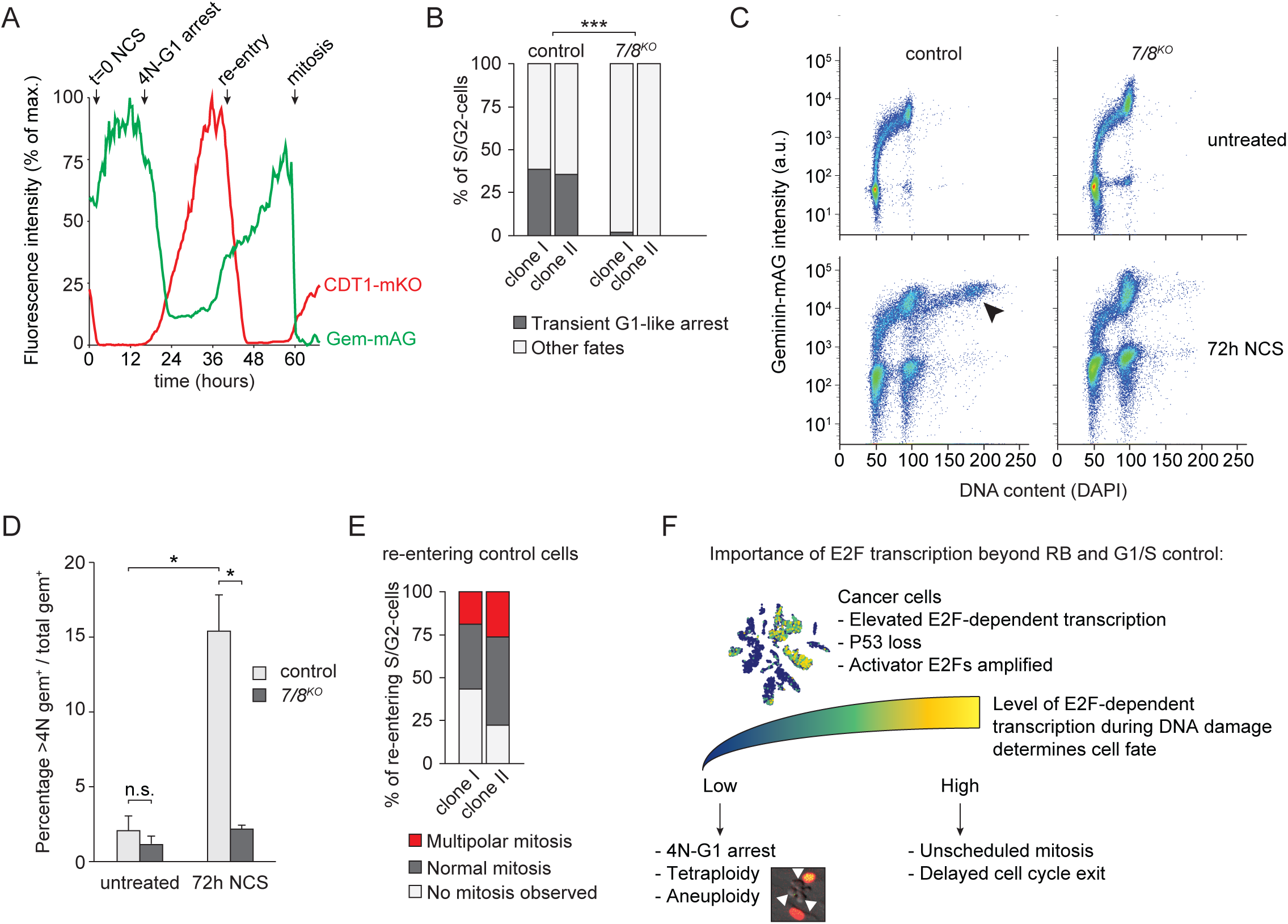
Endocycles after escape from DNA damage-induced arrest are blocked by E2F7/8 deletion. **A** FUCCI fluorescence intensity over time in one representative control cell that underwent a 4N-G1 arrest and subsequently re-entered the cell cycle after ∼2 days. Mitosis was confirmed by inspecting the differential interference contrast (DIC) images. **B** Stacked bar graphs showing the percentages of S/G2 cells that re-entered the cell cycle after a transient 4N-G1 arrest, seen by the re-emergence of Geminin-mAG and disappearance of CDT1-mKO. Cell fates were determined by live cell imaging over a period of 60 hours. Per clone, n=100 cells were followed. *** *P*<0.005. **C** Flow cytometry data showing DNA content and Geminin-mAG expression in RPE cells before and 72 hours after treatment with 100 ng/mL NCS. Arrowhead; population of 8N tetraploid G2 cells. **D** Quantification of percentage >4N Geminin-positive cells, measured by flow cytometry. Bars represent the average ± s.e.m. of in total 4 replicates. *; *P*<0.01. **E** Quantification of the cell cycle fates of control cells that re-entered the cell cycle after a transient 4N-G1 arrest upon treatment with 100 ng/ml NCS. Per clone, n=50 cells were followed. **F** Schematic overview summarizing the main findings of the current study.

## Discussion

High levels of E2F-dependent dependent transcription are seen in most cancers, and correlate with poor prognosis. This can be explained by the fact that it corresponds with increased numbers of cycling cells. However, we now show that E2F-dependent transcription can also be abnormally high in single cycling cancer cells, which has a profound impact on their DNA damage responses and cell cycle fate decision-making (Figure 6F). We studied this impact mechanistically by utilizing the unique feature of atypical E2Fs that they can strongly repress E2F-dependent transcription during S- and G2-phase, without inhibiting the G1/S checkpoint. We provide evidence that uncontrolled E2F-dependent transcription precludes a cell cycle exit and thus allows cells to continue cycling despite severe DNA damage.

Furthermore our data show that this heterogeneity in E2F-dependent transcription during the induction of DNA breaks affects the decision of cells to undergo tetraploidization.

Our mechanistic studies are largely in line with previous studies showing that a P53 can arrest the cell cycle via activation of APC/C^Cdh1^ in response to genotoxic stress. The P53 target P21 inhibits Emi1, which would keep APC/C^Cdh1^ inactive until mitosis under unperturbed conditions. Through this mechanism of APC/C^Cdh1^ activation during G2-phase, cells can degrade and transcriptionally silence multiple cell cycle proteins, to allow cells to enter a state of senescence (Lee, J., Kim, et al, 2009; Wiebusch, and Hagemeier, 2010). The current work shows that P21 is not sufficient to enforce this arrest, because atypical repressor E2Fs also play a critical role. Our results raise the question why mammalian cells would require both atypical E2Fs and P53-P21 to control Emi1 expression and the subsequent 4N-G1 arrest in response to DNA damage. The most logical reason is that this cell cycle arrest is so important to maintain genomic integrity that multiple routes back each other up.

Multiple mechanisms could explain how DNA damage can activate E2F7/8 to mediate the cell cycle arrest after completion of S-phase. First, E2F7 is a direct transcriptional target of P53, meaning that E2F7 can act in parallel with P21, downstream of P53 (Aksoy, Chicas, et al, 2012; Carvajal, Hamard, et al, 2012). Secondly, we recently identified cyclin F-as negative regulator of atypical E2Fs during G2 of unperturbed cell cycles (Yuan, Liu, et al, 2019). Work from the Pagano lab showed that cyclin F is inactivated via ATM in response to genotoxic damage (D’Angiolella, Donato, et al, 2012). This cyclin F-inactivation would then lead to reduced degradation of E2F7/8. Together these P53- and cyclin F-dependent mechanisms could explain why E2F7/8 activity can accumulate in G2 cells after DNA damage to mediate a cell cycle arrest.

Another major finding of the present study was that clearly not all arrested cells become senescent after DNA damage; many can escape this arrest to become tetraploid. Induction of senescence prevents proliferation of cells with damaged DNA (Baus, Gire, et al, 2003; Krenning, Feringa, et al, 2014). Thus, this exit is believed to be an important first line of defense against tumor formation (Bartkova, Horejsi, et al, 2005; Gire, and Dulic, 2015). Stochastic oscillations in P53 could induce an escape from such an arrest resulting in tetraploidy (Reyes, 2018).. Tetraploidization precedes malignant transformation In various cancer types, and therefore this escape could potentially lead to oncogenic transformation (Davoli, and de Lange, 2011; Tanaka, Goto, et al, 2018). Tetraploidy is associated with the ability of non-transformed cells to form tumors in xenografted mice (Davoli, and de Lange, 2012). In addition, tetraploidization may affect drug sensitivity of cells (Lee, A. J., Endesfelder, et al, 2011).

Importantly, *E2F7/8*^*KO*^ cells with elevated E2F transcription failed to become polyploid. Taking this into consideration, one could reason that prevention of tetraploidization by abnormally high E2F-transcription could have a tumor-suppressing effect. However, our work shows that uncontrolled E2F target gene expression during S/G2-phase profoundly increased the number of unscheduled mitoses under DNA damaging conditions. Unscheduled mitosis of diploid cells after DNA damage could be even more detrimental than appearance of tetraploid cells. In the liver for example tetraploidization prevents formation of hepatocellular carcinoma, presumably because polyploid cells have a decreased risk to suffer loss of important tumor suppressor genes (Zhang, Zhou, et al, 2018). Liver-specific deletion of E2f7/8 prevented polyploidization and promoted liver tumor growth, further supporting this notion (Chen, Ouseph, et al, 2012; Kent, Rakijas, et al, 2016; Pandit, Westendorp, et al, 2012).

Although E2F7/8 and Emi1 are rarely mutated in cancer, we show now using single cell transcriptomic data from human cancer patients that E2F-dependent transcription can be strongly induced in cycling malignant cells. We envision that such induction during S- and G2-phase could for example occur via amplification of the activating E2F family members *E2F1, E2F3*, or via enhanced upstream transcription via *MYC*. This would then provide an important mechanism for cells to prevent undergoing a 4N-G1 arrest in response to DNA damaging drugs. It should be noted that tumor cells with elevated E2F transcription would need to activate DNA repair mechanisms in order to prevent excessive DNA damage. Interestingly, many DNA repair pathways, such as mismatch repair, base excision repair, and homologous recombination are all transcriptionally controlled by E2Fs. Future research will need to show which molecular mechanisms can establish heterogeneity in E2F-transcription in cycling cancer cells, and whether this plays a role in resistance to radiation or chemotherapeutic drugs.

## Methods

### Cell culture, cell line generation and transfection

RPE1-hTERT and HEK 293T cell lines were purchased from ATCC and cultured in DMEM (41966052; Thermo Fisher Scientific) containing 10% fetal bovine serum (10500064; Thermo Fisher Scientific). Neocarzinostatin was purchased from Sigma-Aldrich (N9162).

The strategy to produce E2F7/8 double mutant RPE cell lines and confirmation of deletion are described in (Yuan, Liu, et al, 2019). Briefly, cells were transduced with a lentiviral expression vector encoding both Flag-tagged Cas9 and a single guide (sg) RNA sequence against E2F7 (sgRNA #1: GTGCTGCCAGCCCAGATATA, sgRNA #2: GAGCTAGAAACTTCTGGCAC) or E2F8 (sgRNA #1: GTTCCTCTGCCACTTCGTCA, sgRNA #2: GATCTCTGTTGCGGATCTCA). Lentiviral particles were produced by co-transfecting the constructs with third-generation packaging plasmids into HEK 239T cells. The sgE2F7 and sgE2F8 vectors contained puromycin and blasticidin resistance cassettes, respectively, thus allowing for sequential selection of E2F7 and E2F8 mutant clones by manual picking. Cells expressing the vector containing only Cas9, but no sgRNA, served as control cell lines.

The lentiviral construct containing a truncated version of 53BP1 tagged with mApple was a gift from Ralph Weissleder (Addgene plasmid # 69531; http://n2t.net/addgene:69531; RRID:Addgene_69531)), fluoresecent tag was changed to mTurquoise2 using Gibson Assembly.

Lentivirus was produced by transfecting HEK 293T cells with 10 μg lentiviral packaging plasmids (1:1:1) and 10 μg of the desired construct with PEI for 2 h. Then, 10 ml fresh medium was added and virus was harvested after 48 h. Three milliliters of virus containing medium and polybrene (8 μg/ml) was added to RPE cells for an incubation of 24 h. RPE cells containing the construct were selected with puromycin (1.0 μg/ml) for 5 days.

For siRNA experiments, RPE cells were transfected, unless stated otherwise, with a final concentration of 10nM siRNA and 2uL/ml RNAiMAX reagent used according to the manufacturers’ instructions (Life Technologies, 13778030). The following siRNAs were used: Dharmacon D-001210-02-05 (Scrambled), LQ-003329-00-0002 (siTP53), LQ-003296-02-0002 (siRB1), and L-012434-00-0020 (siFBXO5).

### Microscopy

For live cell imaging, 1000 RPE cells were seeded into a 8-well μ-slide (Ibidi). Images were acquired every 20 minutes for 72 hours on a Nikon A1R-STORM microscope using a 10x objective in a humidified chamber at 37°C, 5% CO_2_. Cell tracking was performed manually. Subsequently, fluorescent intensity of CDT1-mKusabira Orange (mKO) and Geminin-mAzami Green (mAG) in these ROIs (region of interest) was automatically measured using NIS Elements software. A cell was considered Geminin or CDT1 positive if the signal in three consecutive frames was higher than 10% of the maximum. Cell cycle phases were assigned, G1-phase: CDT1 positive, early S-phase: CDT1 and Geminin positive and late S/G2-phase: Geminin positive. Mitosis was determined by nuclear envelop breakdown and observation of cell division using the DIC (differential interference contrast) image.

VyCAP microscopic images were quantified with an in-house built ImageJ macro. Images of each punched cell, stored in folders labeled according to plateID and well ID are recursively quantified by creating a ROI in the Hoechst image, and subsequent measurement of FUCCI fluorescence (cell cycle) and DiD fluorescence (*E2F7/8*^*KO*^ vs. control). The outputs of these quantifications were then merged in Rstudio to create a metadata file.

Images for 53BP1 foci analysis were acquired on a Ti-E microscope (Nikon, Japan). ROIs were created using an overlay of the CDT1-mKO and Geminin-mAG channel using an in-house built ImageJ macro. Fluorescent intensity of CDT1-mKO and Geminin-mAG and 53BP1 foci per ROI were analyzed in a semi-automated manner in ImageJ. A cell was considered Geminin or CDT1 positive if the signal was higher than 10% of the maximum.

For immunofluorescence staining, cells were seeded on coverslips. Prior to fixation cells were treated for 1 minute with 0.2% ice-cold Triton X-100. Then cells were fixed with 4% PFA for 20 minutes and permeabilized with 0.1% Trition X-100 for 1 minute at room temperature. Cells were blocked using 5% goat serum and treated with primary antibodies, secondary antibodies and DAPI (Sigma-Aldrich). Subsequently, coverslips were mounted and analyzed on a Leica SPE-II (DMI4000) microscope. For each image ROIs were determined based on DAPI signal and γH2AX foci per cell were quantified using the Find Maxima function in a custom made ImageJ macro. At least 4 different images and 250 cells per condition were analyzed.

### Flow cytometry

For FACS sorting experiments cells were resuspended in DMEM. Sorting was performed based on Geminin-mAG and CDT1-mKO2 signal on a BD-influx. For RNA analysis 50.000 per condition were collected in RLT-buffer. For clonogenic survival assays, cells were collected in complete cell culture medium.

For DNA content and fluorescent intensity analysis, cells were harvested by trypsinization and subsequent fixation with 70% ethanol and overnight storage at 4°C. Before staining, cells were washed twice with ice-cold Tris-buffered saline (TBS) and re-suspended in 500 μl staining buffer that contained 0.25μg/ml DAPI (F6057; Sigma-Aldrich), 250 μg/ml RNase A (RNASEA-RO; Roche), and 0.1% bovine serum albumin (A8531; Sigma-Aldrich). Samples were loaded on a BD FACS Canto II flow cytometer.

Quantification of ploidy and Geminin positivity were conducted using FlowJo v10.0 software.

### Single cell RNA-sequencing

For each experiment, either control or E2F7/8KO cells were labelled with DiD Vybrant dye (ThermoFisher), to distinguish between those two genotypes. After trypsinization, mutant and control cells were resuspended 1:1 in 1 mL of serum-free DMEM containing 1:20000 Hoechst33342 dye (ThermoFisher), and loaded on a VyCAP chip (VyCAP B.V. The Netherlands). The entire chip was scanned on a Ti-E microscope (Nikon, Japan) using the following filter cubes from Chroma; 49000 (Hoechst 33342), 49002 (Geminin-mAG), 49004 (CDT1-mKO), 49006 (DiD).

Using custom VyCAP software we then randomly selected wells containing one cell (detected by Hoechst signal) to punch into 384-well plates. These plates contain 100 nL of barcoded CEL-Seq2 primers + dNTPS, which was protected from evaporation by 5 μL mineral oil. After punching, the plates were covered with adhesive foil, centrifuged for 1 minute at 2500 G, and immediately stored at −80° until further processing.

The library preparations, Illumina sequencing, and read alignment to the human genome (Hg19) were performed by Single Cell Discoveries as described before (Muraro, Dharmadhikari, et al, 2016). The mapped reads were counted using STAR (Dobin, Davis, et al, 2013) and then further processed in Rstudio (version 1.2.5019, Rstudio Team, 2019) and R (version 3.6.1, R Core Team, 2019) using the packages Seurat (Butler, Hoffman, et al, 2018). Cells expressing <1000 or>10000 genes, <4000 Unique Molecular Identifier (UMI) counts, and >30 percent mitochondrial genes where filtered out. Next, data where normalized and scaled with the SCTransform method (Hafemeister, and Satija, 2019), and principle component analysis was used for dimension reduction. Based on the elbow plot, only the first six principle components where used for the clustering analysis. Clusters were visualized with t-Distributed Stochastic Neighbor Embedding (t-SNE). Differentially expressed genes between clusters were obtained using a parametric Wilcoxon rank sum test at a significance level of *P*<0.05. Additionally, pseudo time analysis was performed with the Monocle package (Trapnell, Cacchiarelli, et al, 2014).

### Analysis of public available single cell transcriptomics datasets

Normalized count data and metadata from studies GSE103322 (head and neck squamous cell carcinoma) and GSE116256 (acute myeloid leukemia) were downloaded from Gene Expression Omnibus. Data were processed using the Seurat package as described above. E2F scores were obtained by calculating mean z-scores for a panel of 80 E2F target genes. The methodology to determine cell type identities, classification into malignant versus normal cells, and classification into cycling versus non-cycling cells are described in detail in the original publications (Puram, Tirosh, et al, 2017; van Galen, Hovestadt, et al, 2019). From study GSE116256 we included only samples from patients prior to treatment, and with unambiguous malignancy classifications (12 out of 16). Statistical analysis of E2F scores between various normal and malignant cells was performed using a Kruskall-Wallis test followed by Dunn’s test of multiple comparisons, with Benjamini-Hochberg post-hoc *P*-value correction.

### Quantitative PCR

Isolation of RNA, cDNA synthesis, and quantitative PCR were performed based on the manufacturers’ instructions for QIAGEN (RNeasy Kits), Thermo Fisher Scientific (cDNA synthesis Kits), and Bio-Rad (SYBR Green Master Mix), respectively. Gene transcript levels were determined using the ΔΔCt method for multiple-reference gene correction. *GAPDH* and *RSP18* were used as reference genes for normalization. qPCR primer sequences are provided in Table EV1.

### Colony formation assays

For colony formation assays cells were, if indicated, pretreated with NCS. Geminin positive cells were collected using FACS sorting as described and plated at low density (250 cells/well) in a 6 well plate. Medium was replenished every 48 hours and cells were fixed after 10 days. Fixation was performed with Acitic acid:Methanol (1:7 vol/vol) for 5 minutes, then cells were washed with PBS and incubated with Crystal Violet staining solution (0.5% Crystal Violet, Sigma-Aldrich) for 2 hours at room temperature.Subsequently dishes were rinsed with water and air-dried. Number of colonies per well was counted manually, each condition was performed in duplo.

### Statistics

Immunoblot, flow cytometry, and qPCR results were repeated four times unless otherwise described in the figure legends. Statistical analyses on qPCR was done by a Student’s t-test for figure 2A and a Kruskal-Wallis test with a post hoc Dunnett’s test for figure 5D and S5C,E. Cumulative curves from Figs 2B, 2F and 2H were analyzed by log-rank tests. * P < 0.05, ** P < 0.005, ***P <0.001 unless indicated otherwise.

## Data availablity

Raw and processed single cell RNA-sequencing data generated in this study are available on Gene Expression Omnibus under accession number GSE146759.

## Acknowledgements

We thank Judith Vivié and Mauro Muraro (Single Cell Discoveries, NL) for support with single cell sequencing library preparation and sequencing services. Ger Arkesteijn (Faculty of Veterinary Medicine, Utrecht University, NL), Reinier van der Linden and Stefan van der Elst (Hubrecht Institute-KNAW, NL) for assistance with FACS sorting experiments. This work is financially supported by the China Scholarship Council (CSC) (files no. 201306380101 to RY); by KWF Kankerbestrijding (Dutch Cancer Society project grants UU2013-5777 and 11941-2018-II); and by a ZonMW (grant 91116011). Further financial support was provided by a research infrastructure grant by Utrecht Life Sciences.

## Author contributions

H.A.S. conceived and performed experiments, analyzed data, and wrote the manuscript. L.M.v.R., E.M., and Y.R. conceived, analyzed, and performed experiments and analyzed data. F.M.R. performed bioinformatic analysis. R.W. provided expert support with live cell imaging experiments and data analysis. A.d.B. provided mentorship and wrote the manuscript. B.W. conceived and oversaw the study, analyzed data, and wrote the manuscript.

## Reference List

Aksoy, O., Chicas, A., Zeng, T., Zhao, Z., McCurrach, M., Wang, X., and Lowe, S.W. (2012). The atypical E2F family member E2F7 couples the p53 and RB pathways during cellular senescence. Genes Dev. 14, 1546–1557.

Arora, M., Moser, J., Phadke, H., Basha, A.A., and Spencer, S.L. (2017). Endogenous Replication Stress in Mother Cells Leads to Quiescence of Daughter Cells. Cell. Rep. 7, 1351–1364.

Bartkova, J., Horejsi, Z., Koed, K., Kramer, A., Tort, F., Zieger, K., Guldberg, P., Sehested, M., Nesland, J.M., Lukas, C. et al. (2005). DNA damage response as a candidate anti-cancer barrier in early human tumorigenesis. Nature 7035, 864–870.

Bassermann, F., Frescas, D., Guardavaccaro, D., Busino, L., Peschiaroli, A., and Pagano, M. (2008). The Cdc14B-Cdh1-Plk1 axis controls the G2 DNA-damage-response checkpoint. Cell 2, 256–267.

Baus, F., Gire, V., Fisher, D., Piette, J., and Dulic, V. (2003). Permanent cell cycle exit in G2 phase after DNA damage in normal human fibroblasts. EMBO J. 15, 3992–4002.

Bertoli, C., Herlihy, A.E., Pennycook, B.R., Kriston-Vizi, J., and de Bruin, R.A. (2016). Sustained E2F-Dependent Transcription Is a Key Mechanism to Prevent Replication-Stress-Induced DNA Damage. Cell. Rep. 7, 1412–1422.

Bertoli, C., Skotheim, J.M., and de Bruin, R.A. (2013). Control of cell cycle transcription during G1 and S phases. Nat. Rev. Mol. Cell Biol. 8, 518–528.

Boekhout, M., Yuan, R., Wondergem, A.P., Segeren, H.A., van Liere, E.A., Awol, N., Jansen, I., Wolthuis, R.M., de Bruin, A., and Westendorp, B. (2016). Feedback regulation between atypical E2Fs and APC/CCdh1 coordinates cell cycle progression. EMBO Rep. 3, 414–427.

Butler, A., Hoffman, P., Smibert, P., Papalexi, E., and Satija, R. (2018). Integrating single-cell transcriptomic data across different conditions, technologies, and species. Nat. Biotechnol. 5, 411–420.

Cappell, S.D., Chung, M., Jaimovich, A., Spencer, S.L., and Meyer, T. (2016). Irreversible APC(Cdh1) Inactivation Underlies the Point of No Return for Cell-Cycle Entry. Cell 1, 167–180.

Carvajal, L.A., Hamard, P.J., Tonnessen, C., and Manfredi, J.J. (2012). E2F7, a novel target, is up-regulated by p53 and mediates DNA damage-dependent transcriptional repression. Genes Dev. 14, 1533–1545.

Chao, H.X., Poovey, C.E., Privette, A.A., Grant, G.D., Chao, H.Y., Cook, J.G., and Purvis, J.E. (2017). Orchestration of DNA Damage Checkpoint Dynamics across the Human Cell Cycle. Cell. Syst. 5, 445-459.e5.

Chen, H.Z., Ouseph, M.M., Li, J., Pecot, T., Chokshi, V., Kent, L., Bae, S., Byrne, M., Duran, C., Comstock, G. et al. (2012). Canonical and atypical E2Fs regulate the mammalian endocycle. Nat. Cell Biol. 11, 1192–1202.

D’Angiolella, V., Donato, V., Forrester, F.M., Jeong, Y.T., Pellacani, C., Kudo, Y., Saraf, A., Florens, L., Washburn, M.P., and Pagano, M. (2012). Cyclin F-mediated degradation of ribonucleotide reductase M2 controls genome integrity and DNA repair. Cell 5, 1023–1034.

Davoli, T., and de Lange, T. (2012). Telomere-driven tetraploidization occurs in human cells undergoing crisis and promotes transformation of mouse cells. Cancer cell 6, 765–776.

Davoli, T., and de Lange, T. (2011). The causes and consequences of polyploidy in normal development and cancer. Annu. Rev. Cell Dev. Biol. 585–610.

Dobin, A., Davis, C.A., Schlesinger, F., Drenkow, J., Zaleski, C., Jha, S., Batut, P., Chaisson, M., and Gingeras, T.R. (2013). STAR: ultrafast universal RNA-seq aligner. Bioinformatics 1, 15–21.

Gire, V., and Dulic, V. (2015). Senescence from G2 arrest, revisited. Cell. Cycle 3, 297–304.

Hafemeister, C., and Satija, R. (2019). Normalization and variance stabilization of single-cell RNA-seq data using regularized negative binomial regression. bioRxiv 576827.

Helmbold, H., Kömm, N., Deppert, W., and Bohn, W. (2009). Rb2/p130 is the dominating pocket protein in the p53-p21 DNA damage response pathway leading to senescence. Oncogene 39, 3456–3467.

Kent, L.N., and Leone, G. (2019). The broken cycle: E2F dysfunction in cancer. Nat. Rev. Cancer. 6, 326–338.

Kent, L.N., Rakijas, J.B., Pandit, S.K., Westendorp, B., Chen, H.Z., Huntington, J.T., Tang, X., Bae, S., Srivastava, A., Senapati, S. et al. (2016). E2f8 mediates tumor suppression in postnatal liver development. J. Clin. Invest. 8, 2955–2969.

Krenning, L., Feringa, F.M., Shaltiel, I.A., van den Berg, J., and Medema, R.H. (2014). Transient activation of p53 in G2 phase is sufficient to induce senescence. Mol. Cell 1, 59–72.

Lan, W., Bian, B., Xia, Y., Dou, S., Gayet, O., Bigonnet, M., Santofimia-Castano, P., Cong, M., Peng, L., Dusetti, N., and Iovanna, J. (2018). E2F signature is predictive for the pancreatic adenocarcinoma clinical outcome and sensitivity to E2F inhibitors, but not for the response to cytotoxic-based treatments. Sci. Rep. 1, 8330–z.

Lee, A.J., Endesfelder, D., Rowan, A.J., Walther, A., Birkbak, N.J., Futreal, P.A., Downward, J., Szallasi, Z., Tomlinson, I.P., Howell, M., Kschischo, M., and Swanton, C. (2011). Chromosomal instability confers intrinsic multidrug resistance. Cancer Res. 5, 1858–1870.

Lee, J., Kim, J.A., Barbier, V., Fotedar, A., and Fotedar, R. (2009). DNA damage triggers p21WAF1-dependent Emi1 down-regulation that maintains G2 arrest. Mol. Biol. Cell 7, 1891–1902.

Li, J., Ran, C., Li, E., Gordon, F., Comstock, G., Siddiqui, H., Cleghorn, W., Chen, H.Z., Kornacker, K., Liu, C.G. et al. (2008). Synergistic function of E2F7 and E2F8 is essential for cell survival and embryonic development. Dev. Cell. 1, 62–75.

Muraro, M.J., Dharmadhikari, G., Grun, D., Groen, N., Dielen, T., Jansen, E., van Gurp, L., Engelse, M.A., Carlotti, F., de Koning, E.J., and van Oudenaarden, A. (2016). A Single-Cell Transcriptome Atlas of the Human Pancreas. Cell. Syst. 4, 385-394.e3.

Pandit, S.K., Westendorp, B., Nantasanti, S., van Liere, E., Tooten, P.C., Cornelissen, P.W., Toussaint, M.J., Lamers, W.H., and de Bruin, A. (2012). E2F8 is essential for polyploidization in mammalian cells. Nat. Cell Biol.

Puram, S.V., Tirosh, I., Parikh, A.S., Patel, A.P., Yizhak, K., Gillespie, S., Rodman, C., Luo, C.L., Mroz, E.A., Emerick, K.S. et al. (2017). Single-Cell Transcriptomic Analysis of Primary and Metastatic Tumor Ecosystems in Head and Neck Cancer. Cell 7, 1611-1624.e24.

Sakaue-Sawano, A., Kurokawa, H., Morimura, T., Hanyu, A., Hama, H., Osawa, H., Kashiwagi, S., Fukami, K., Miyata, T., Miyoshi, H. et al. (2008). Visualizing spatiotemporal dynamics of multicellular cell-cycle progression. Cell 3, 487–498.

Stevens, M., Oomens, L., Broekmaat, J., Weersink, J., Abali, F., Swennenhuis, J., and Tibbe, A. (2018). VyCAP’s puncher technology for single cell identification, isolation, and analysis. Cytometry A. 12, 1255–1259.

Tanaka, K., Goto, H., Nishimura, Y., Kasahara, K., Mizoguchi, A., and Inagaki, M. (2018). Tetraploidy in cancer and its possible link to aging. Cancer science 9, 2632–2640.

Thurlings, I., Martinez-Lopez, L.M., Westendorp, B., Zijp, M., Kuiper, R., Tooten, P., Kent, L.N., Leone, G., Vos, H.J., Burgering, B., and de Bruin, A. (2017). Synergistic functions of E2F7 and E2F8 are critical to suppress stress-induced skin cancer. Oncogene 6, 829–839.

Trapnell, C., Cacchiarelli, D., Grimsby, J., Pokharel, P., Li, S., Morse, M., Lennon, N.J., Livak, K.J., Mikkelsen, T.S., and Rinn, J.L. (2014). The dynamics and regulators of cell fate decisions are revealed by pseudotemporal ordering of single cells. Nat. Biotechnol. 4, 381–386.

Vaidyanathan, S., Cato, K., Tang, L., Pavey, S., Haass, N.K., Gabrielli, B.G., and Duijf, P.H. (2016). In vivo overexpression of Emi1 promotes chromosome instability and tumorigenesis. Oncogene 41, 5446–5455.

van Galen, P., Hovestadt, V., Wadsworth Ii, M.H., Hughes, T.K., Griffin, G.K., Battaglia, S., Verga, J.A., Stephansky, J., Pastika, T.J., Lombardi Story, J. et al. (2019). Single-Cell RNA-Seq Reveals AML Hierarchies Relevant to Disease Progression and Immunity. Cell 6, 1265-1281.e24.

Westendorp, B., Mokry, M., Groot Koerkamp, M.J., Holstege, F.C., Cuppen, E., and de Bruin, A. (2012). E2F7 represses a network of oscillating cell cycle genes to control S-phase progression. Nucleic Acids Res. 8, 3511–3523.

Wiebusch, L., and Hagemeier, C. (2010). p53- and p21-dependent premature APC/C-Cdh1 activation in G2 is part of the long-term response to genotoxic stress. Oncogene 24, 3477–3489.

Yuan, R., Liu, Q., Segeren, H.A., Yuniati, L., Guardavaccaro, D., Lebbink, R.J., Westendorp, B., and de Bruin, A. (2019). Cyclin F-dependent degradation of E2F7 is critical for DNA repair and G2-phase progression. EMBO J. e101430.

Zhang, S., Zhou, K., Luo, X., Li, L., Tu, H.C., Sehgal, A., Nguyen, L.H., Zhang, Y., Gopal, P., Tarlow, B.D., Siegwart, D.J., and Zhu, H. (2018). The Polyploid State Plays a Tumor-Suppressive Role in the Liver. Dev. Cell. 4, 447-459.e5.

