## Supplementary Tables and Figures for "Excessive E2F transcription in single cancer cells precludes transient cell cycle exit after DNA damage"

**Table of contents**

1. Supplemental Tables
2. Legends to Supplemental Figures

### 1. Supplementary tables

TABLE EV1: qPCR primers

| | ***Forward primer (5'-3')*** | ***Reverse primer (3'-5')*** |
|---|---|---|
| *GAPDH* | CTCTGCTCCTCCTGTTCG | GCCCAATACGACCAAATCC |
| *TP53* | GTTCCGAGAGCTGAATGAGG | TCTGAGTCAGGCCCTTCTGT |
| *RB1* | GAGACACAAGCAACCTCAGC | GCTCAGACAGAAGGCGTTC |
| *FBXO5* | TGACAGTCTACAATCCTGCCTGC | TTCCTTCAGCATCTCCCGATC |
| *CDC6* | AAACCCGATCCCAGGCACAG | AGGCAGGGCTTTTACACGAGGAG |
| *E2F1* | GACCACCTGATGAATATCTG | TGCTACGAAGGTCCTGAC |
| *CDKN1A* | CTCTAAGGTTGGGCAGGGTGACC | CAGAGGGGGGTATCAAGAGCCAG |
| *CDT1* | CGTCCAGGACATGATGCGTAGG | TTGAAGGTGGGGACACTGCG |
| *MCM2* | GGCAATGATCCTCTCACCTCC | CATCCTCTTCTTCCTCCAGGG |
| *RAD51* | TGCTTATTGTAGACAGTGCCACC | CACCAAACTCATCAGCGAGTC |
| *CCNE1* | GACACCATGAAGGAGGACGG | ATTGTCCCAAGGCTGGCTC |
| *RSP18* | AGTTCCAGCATATTTTGCGAG | CTCTTGGTGAGGTCAATGTC |

TABLE EV2: Antibodies for immunoblots and immunofluorescence staining

| ***Application*** | ***Name*** | ***Company*** | ***Cat #*** | ***Dilution*** |
|---|---|---|---|---|
| Immunoblots | P21 | Santa Cruz | (M-19) sc-471 | 1:1000 |
| | CDC6 | Santa Cruz | Sc-9964 | 1:1000 |
| | RB1 | Santa Cruz | (C-15) sc-50 | 1:1000 |
| | EMI1 | Abcam | ab244426 | 1:1000 |
| | P53 | Calbiochem | OP03 | 1:1000 |
| | γ-tubulin (clone GTU-88) | Sigma | T6557 | 1:1000 |
| Immunofluorescence | γ-H2AX | Cell Signaling | S139 | 1:200 |

### 2. Legends to Supplementary Figures

**Figure S1 – Enhanced E2F-dependent transcription in single tumor cells from patients with acute myeloid leukemia (AML).**

E2F target gene expression scores in malignant versus non-malignant cells from 10 different AML patients, prior to chemotherapy treatment. Cells are color-labeled according to binary cell cycle classification scores, as determined in the original publication (van Galen, Hovestadt, et al, 2019)].

**Figure S2 – Clustering analysis of malignant and non-malignant cells from patients with acute myeloid leukemia (AML).**

**A** Dimensionality reduction using t-Stochastic Neighbourhood Embedding (tSNE), labelled by malignancy classification. Malignant cells were distinguished from non-malignant cells according to a machine-learning algorithm described in the original publication (van Galen, Hovestadt, et al, 2019).

**B** Same tSNE map as in A, but now labeled according to patient ID.

**C** Same tSNE map as in B, but now labeled according to cell-type, as determined in the original publication (van Galen, Hovestadt, et al, 2019).

**Figure S3 – Generation and characterization of *E2F7/8$^{KO}$* RPE-FUCCI cell lines.**

**A** Schematic representation of the FUCCI system, based on alternating expression of fluorescent-tagged truncated CDT1 and geminin.

**B** Schematic overview of experimental strategy to create *E2F7/8$^{KO}$* cells after CRISPR/Cas9.

**C** Schematic representation of live cell imaging to monitor two possible fates of RPE-FUCCI cells in G1-phase during, and quantification of fractions of unperturbed control and *E2F7/8$^{KO}$* cells that remained in G1 for the entire duration of live cell imaging. Two independently created CRISPR clones were analyzed, n=200 cells per condition.

**D** Cell proliferation, measured by total cell count at indicated time point during the live cell imaging experiment. NCS was added at t=0 hours. Data indicate mean ± standard deviation of two independent clones of mutant or control cells. For each clone, total amount of nuclei in an area of 3x3 stitched images (using a 20x objective ) were counted.

**E** Quantification of 53BP1 foci in G1 cells. Photo shows a representative image of CDT1/53BP1 signals; the right nucleus contains a 53BP1 focus. Bars represent the average of two clones, n=500 cells per condition. Difference was statistically evaluated with a chi-square-test. *** P<0.001

**Figure S4 – Pseudotime alignment of single cell RPE-FUCCI RNA-sequencing data.**

**A** mRNA counts of representative E2F target genes (*CDC6* and *MCM2*) and FOXM1 targets (*CDK1, PLK1*) and the general cell cycle marker *MKI67* from the single cell sequencing data, separated by

FUCCI cell cycle phase and genotype condition, 48 hours after addition of 200 ng/mL NCS. The far majority of the cells were in G1 or 4N-G1.

**B** Monocle pseudotime alignment of all RPE-FUCCI cells, labeled by NCS treatment. The heatmap shows the expression of the main genes explaining the first two components of the data reduction algoritm; selected cell cycle genes (including E2F targets *GINS2*, *RAD51*, *RRM2*, *DBF4*, and *MCM2*) are indicated..

**C** Same Monocle alignment as in B, but separated by NCS treatment and genotype, and color-labeled by cell cycle phase, allocated using measurement of FUCCI fluorescence. Arrowheads refer to Geminin$^{pos}$ E2F7/8$^{KO}$ cells which cluster with G0/1 phase cells (a) and S/G2 phase cells (b).

**Figure S5 – RB does not affect the decision to undergo a 4N-G1 arrest after DNA damage in RPE cells.**

**A** qPCR measurement of *TP53* gene expression, after 48 hours siRNA against TP53 or RB1, in control and *E2F7/8*$^{KO}$ RPE-FUCCI cells. Bars represent mean ± s.e.m. of duplo measurements in two independent cell clones. NCS was added 24 hours prior to harvesting.

**B** Immunoblots showing the effect of TP53 and RB1 RNAi on expression of RB1 protein, 48 hours after transfection, and 24 hours after NCS treatment. Control and *E2F7/8*$^{KO}$ blots were loaded with the same amount of protein, and imaging was done simultaneously, with identical exposure times for each membrane. Gamma-tubulin served as loading control.

**C** qPCR measurement of *RB1* expression after siRNA against TP53 or RB1, in control and *E2F7/8*$^{KO}$ RPE-FUCCI cells. Bars represent mean ± s.e.m. of n=4 biological replicates.

**D** Quantification of the percentages of S/G2 (Geminin-mAG positive) cells 24 and 48 hours after NCS treatment. Scrambled RNAi (scr.) data are the same as in Figure 5B. Per condition at least n=700 cells were counted. Differences in percentages of cells per condition were statistically evaluated using Fisher's Exact tests. * $P< 0.05$, ** $P<0.005$

**E** qPCR measurement of a panel of E2F target genes after siRNA against *TP53* or *RB1*, in control and *E2F7/8*$^{KO}$ RPE-FUCCI cells. Bars represent mean ± s.e.m. of duplo measurements in two independent cell clones. Scrambled RNAi (scr.) data are the same as in Figure 5C.

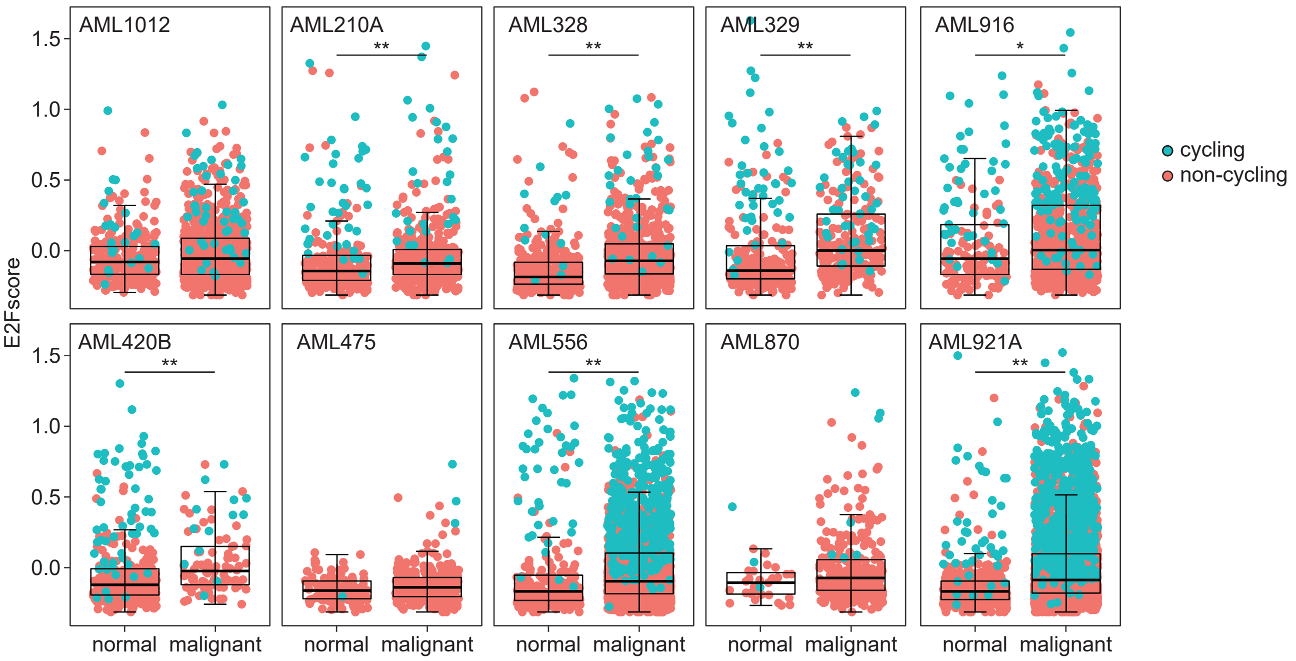


Supplemental Figure S1 - Segeren et al. 2020
Excessive E2F transcription in single cancer cells precludes transient cell cycle exit after DNA damage.

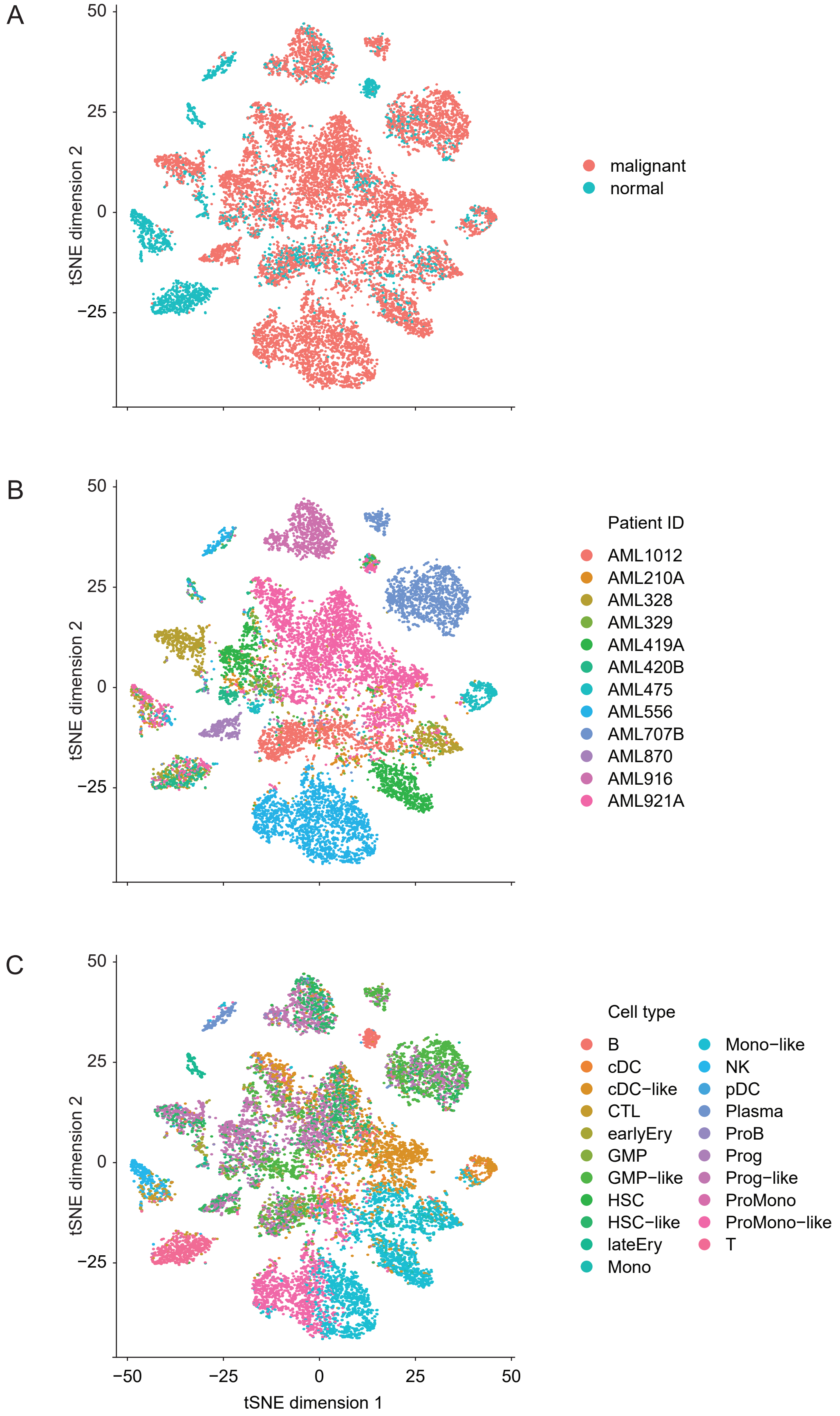


Supplemental Figure S2 - Segeren et al. 2020
Excessive E2F transcription in single cancer cells precludes transient cell cycle exit after DNA damage.

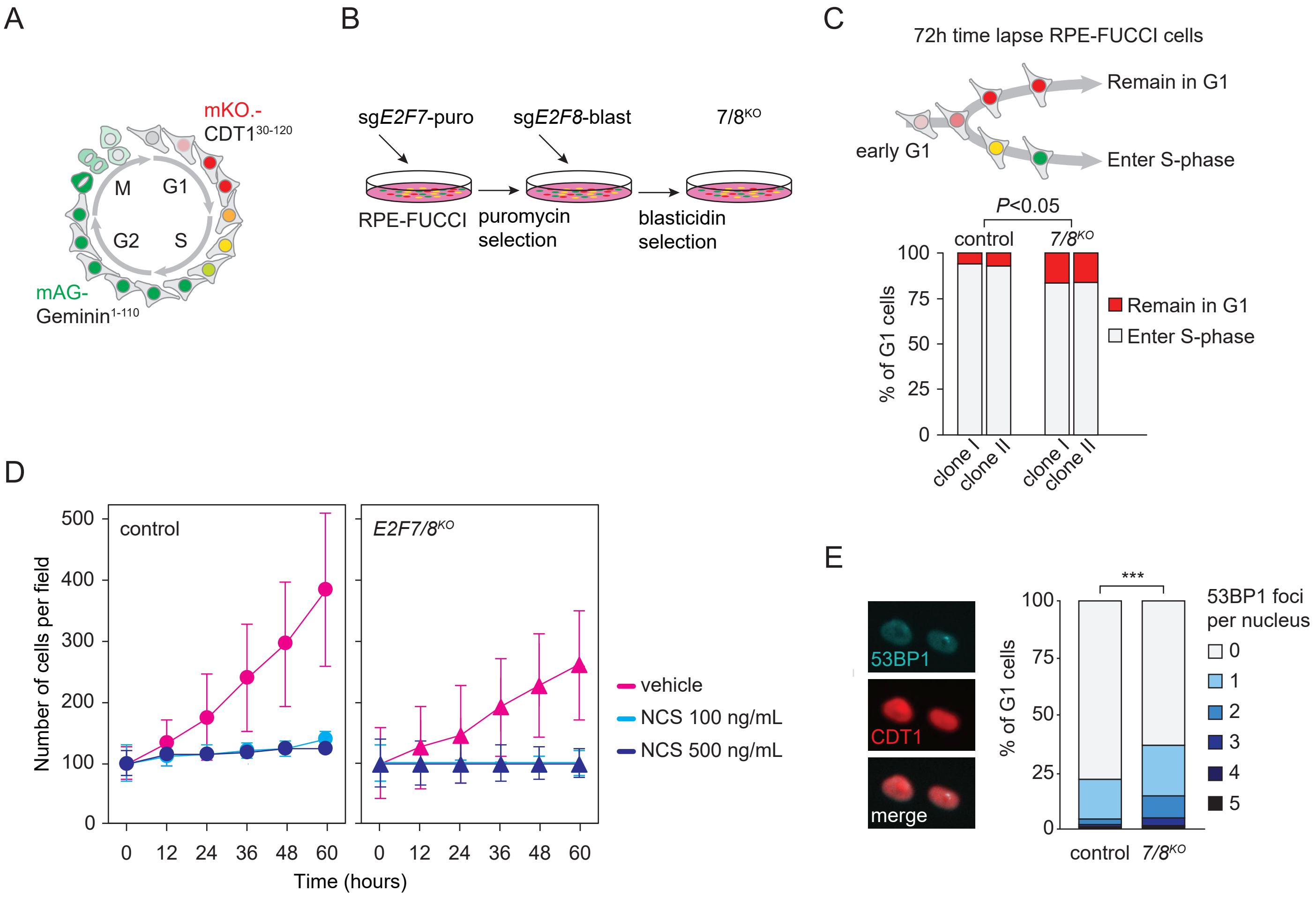


Supplemental Figure S3 - Segeren et al. 2020
Excessive E2F transcription in single cancer cells precludes transient cell cycle exit after DNA damage.

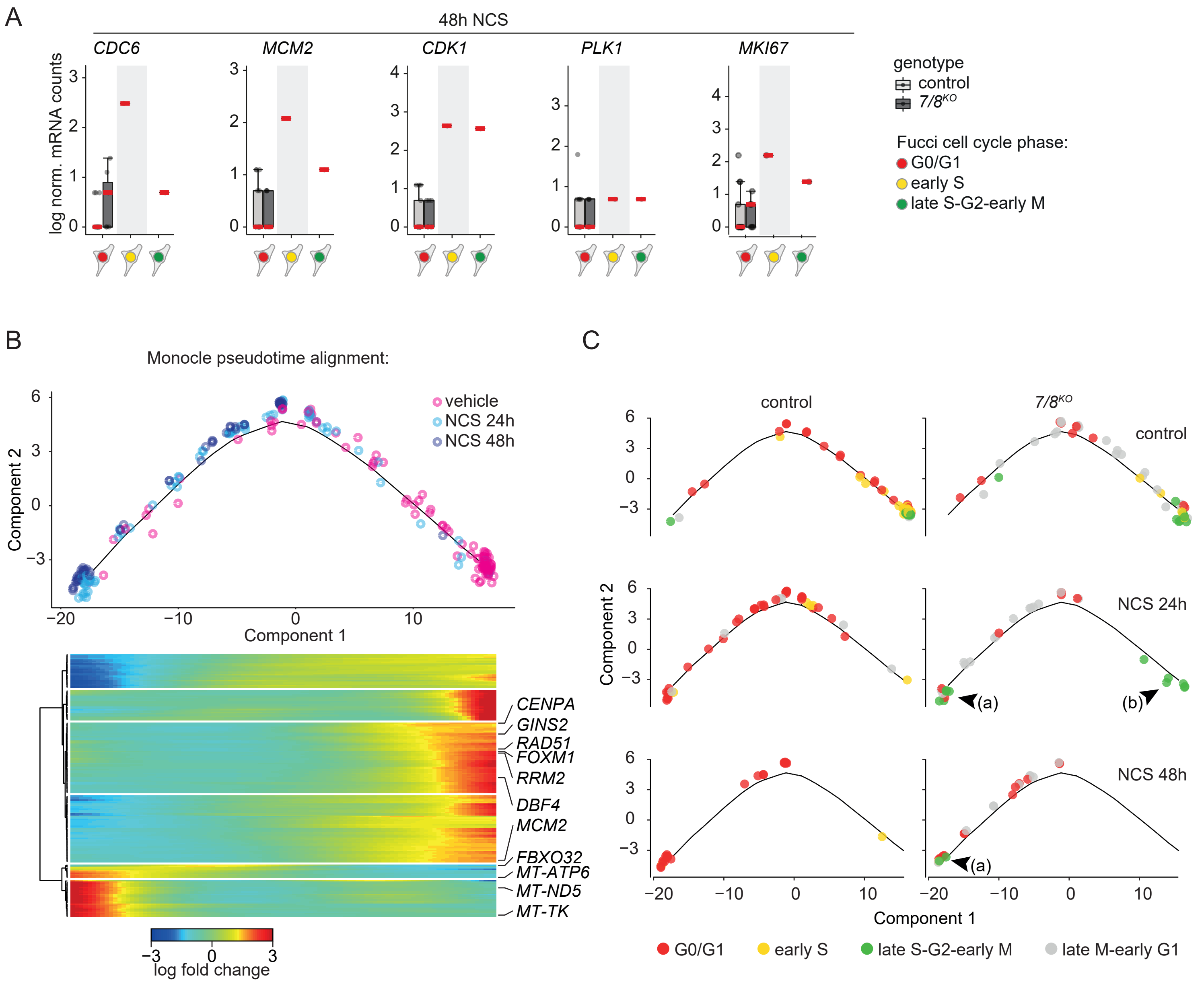


Supplemental Figure S4 - Segeren et al. 2020

Excessive E2F transcription in single cancer cells precludes transient cell cycle exit after DNA damage.

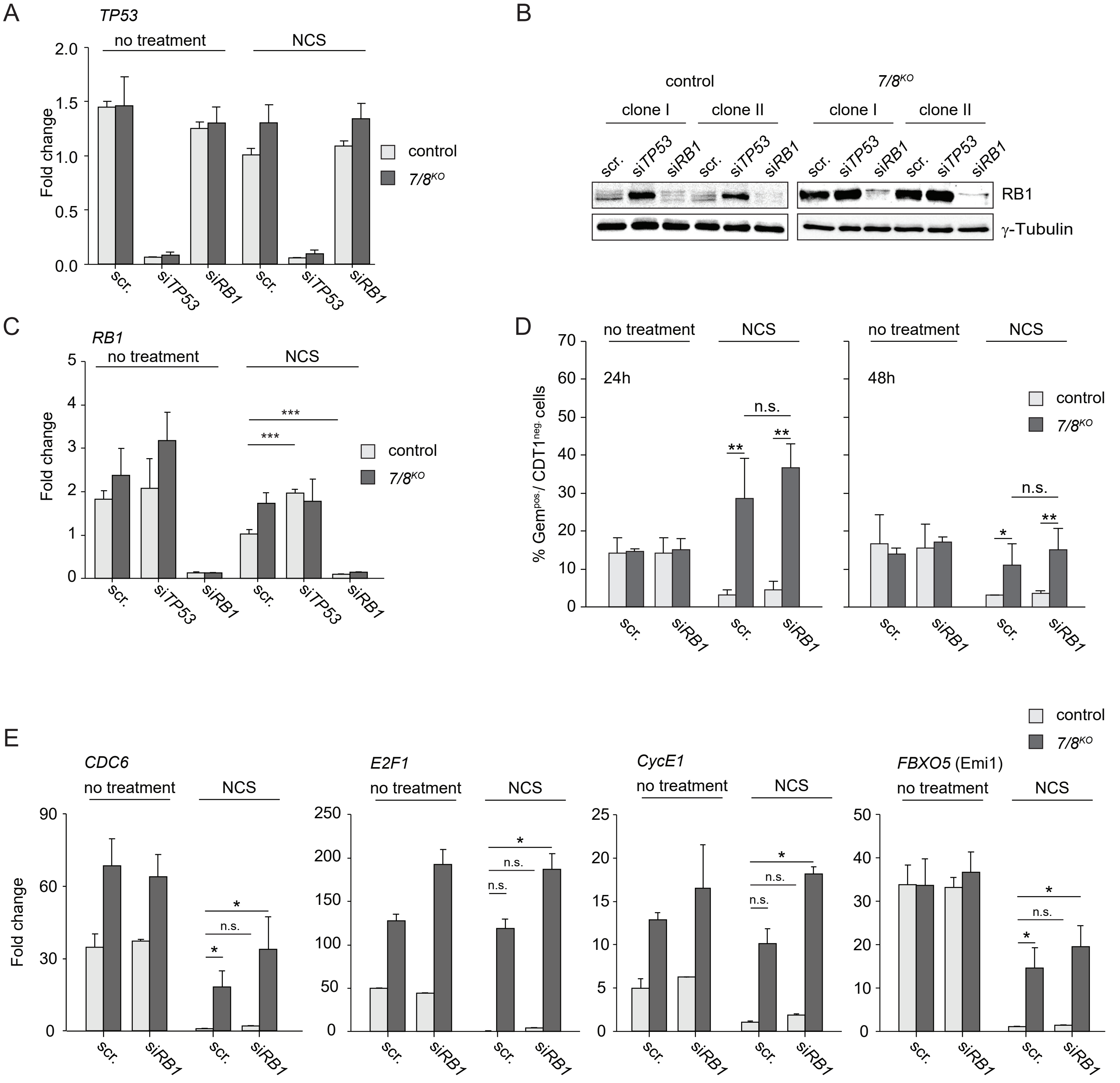


Supplemental Figure S5 - Segeren et al. 2020

Excessive E2F transcription in single cancer cells precludes transient cell cycle exit after DNA damage.
